# Deformation geometry of cellulose fibril arrays constraining the stretching and growth of plant cell walls

**DOI:** 10.64898/2025.12.23.696159

**Authors:** M.C. Jarvis

## Abstract

There are several ways in which the nanoscale array of cellulose fibrils in one layer of a plant cell wall can rearrange to permit the cell wall to expand under external uniaxial tension or, in the case of primary cell-walls, the biaxial turgor pressure that drives growth. Here, seven such deformation modes were identified and their scale-independent geometry was described: fibril rotation, regular shear, interdigitated sliding, fibril stretching, fibril respacing and the formation and straightening of waves. The distinction between regular shear and interdigitated sliding was introduced to capture a continuous range of sliding modes at fibril interfaces.

Combinations of these nanoscale deformations were examined to find out how their relative magnitude must vary to satisfy cell-scale geometric constraints. When the cellulose fibrils were transversely oriented, respacing, wave formation or both were needed for elongation. When the tissue restrained twist to zero, the deformation modes became co-ordinated, readjusting as the cellulose orientation became more axial during elongation. Regular shear, possibly facilitated by expansin activity, could then control the width of the elongating cell-wall. Each deformation mode fortuitously contributed most elongation at the microfibril orientation where it was most efficiently driven by the local force vector.

## 1. Introduction

### 1.1. Background

In multicellular plants, every cell is fixed to the next through the cell walls. The primary cell wall, through its unique capability to stretch without losing strength [1], controls how turgor pressure drives anisotropic growth of each cell [2] in concert with the growth of its neighbours [3]. Cell-wall stretching and synthesis of new wall material are synchronised [4]. From these properties of primary cell walls emerge the growth and diverse forms of living plants [5]. The secondary walls of cells that have ceased to grow provide strength, reinforcing plants against stresses like wind and the negative pressures entailed by xylem translocation [6].

Both primary and secondary cell walls are assembled from arrays of cellulose fibrils embedded in non-cellulosic polymers [1]. There are recent reviews of the polymer structures [1, 7, 8] and new insights are still emerging, e.g. [9, 10]. A key factor in growth and strength is the microfibril angle ϑ between the mean fibril orientation and the cell axis, which results initially from the direction in which the cellulose synthase complexes travel when laying down microfibrils [5]. The initial microfibril angle depends on developmental factors [5, 11] and, it is proposed, by feedback from forces within the cell wall [4, 12, 13].

Structural changes observed under mechanical tension have been described by, e.g. [14, 15]. It is now considered that direct cellulose-cellulose interactions between fibrils are important in the mechanical properties of both primary [15] and secondary [14] cell walls, but these interactions are not continuous along each fibril. Particularly in primary cell walls, cellulose surfaces adhere only in short segments [1], here called *junction zones* [16], forming an anastomosing network.

At the nanometre scale there are various ways in which a small group of fibrils could rearrange to make a local contribution to the expansion of the whole cell wall under stress. It is presumably at this local level in primary cell walls that growth is controlled, although we do not understand the polymer structures and interactions well enough to be sure of the detailed molecular mechanisms responsible [1, 17] and hence the relative energy demands of the possible modes of fibril rearrangement. Both free and activation energies include a mechanical contribution i.e. the product of the local force vector and deformation. While elongation growth has received most attention, accompanying lateral and local expansion also shapes cells and organs.

Similar questions surround the strength of wood as an engineering material [14]. In a landmark paper on the stretching of wood under tension, Keckes et al. [18]. showed how two nanoscale deformation modes, shear between fibrils and rotation of the fibril array, co-operate geometrically to allow elongation. Since then, several other local deformation modes of fibril arrays have been imaged, or inferred from diffraction data, or predicted from coarse-grained simulations; in wood, primary cell walls or both. These deformation modes include fibril stretching [14], widening of the spaces between fibrils [19], forms of sliding different [4, 12, 15, 18] from the regular shear assumed by Keckes et al. [18], and the formation or straightening of waves [20]. The rules defining how all these modes of deformation interact and contribute to cell-wall expansion have not yet been established, although valuable insights have come from coarse-grained simulations [21].

### 1.2. Objectives

The present paper draws on concepts originating in both wood and primary cell walls to arrive at an analytical description of the geometry of each of the known local deformation modes of fibril arrays. Restricting the analysis to geometry allows it to be scale-independent. Thus, nanoscale functions can be scaled up to cell-scale inferences, and it is not a problem that, whereas 3 nm microfibrils are the elementary fibrillar units of primary walls [1], macrofibrils – bundles of microfibrils – form topologically similar anastomosing arrays in secondary walls [14]. In primary cell-walls, each mode of nanoscale deformation is also a point at which there is potential to control the magnitude and direction of growth that results [1]. In wood, each mode of deformation is a link between nanostructure and macroscopic resistance to stress [14]. The aim of this study is to predict the combinations of nanoscale deformation modes needed to generate defined cell-scale changes in dimensions, in scenarios typical of the walls of growing cells and of the deformation of both primary and wood cell walls under external tension.

## 2. Results

### 2.1. Domains of deformation

The functions describing elongation, change in width and twist differed with the shape of the cell-wall domain under consideration. The rectangular domain assumed by [18] corresponds to the cell-wall of a softwood tracheid cut open and laid out flat. However, if the domain is to be a homogeneous rectangle, it seems sensible to consider a single wall facet rather than flattening out all the longitudinal walls. In softwoods the tangential walls are uniform but the radial walls contain regions (pit fields) with different cellulose orientation [22]. In growing tissues the microfibril orientation differs at the cell edges [23]. In many cell types, each wall facet is a rectangle with varying length/width ratio (aspect ratio), although there are also cells with much more complex shapes [24]. The cell edge domains [23] can be examined separately as distinct, narrow rectangles.

If the aim is to examine the deformation of a small domain within a spatially non-uniform cell wall, e.g. an epidermal pavement cell [25], a cell from which a hair is initiating [26], or a pit-field region in a radial wood cell-wall [22], it is more convenient to assume an array of circular domains each deforming to an ellipse. With circular geometry discontinuous functions are avoided, together with other complications arising from twist. The geometry of deformation modes within initially circular domains is described in SI Appendix C.

### 2.2. Individual deformation modes

Four kinds of deformation of an array of straight fibrils are illustrated in Fig.1.

**Fig. 1.**
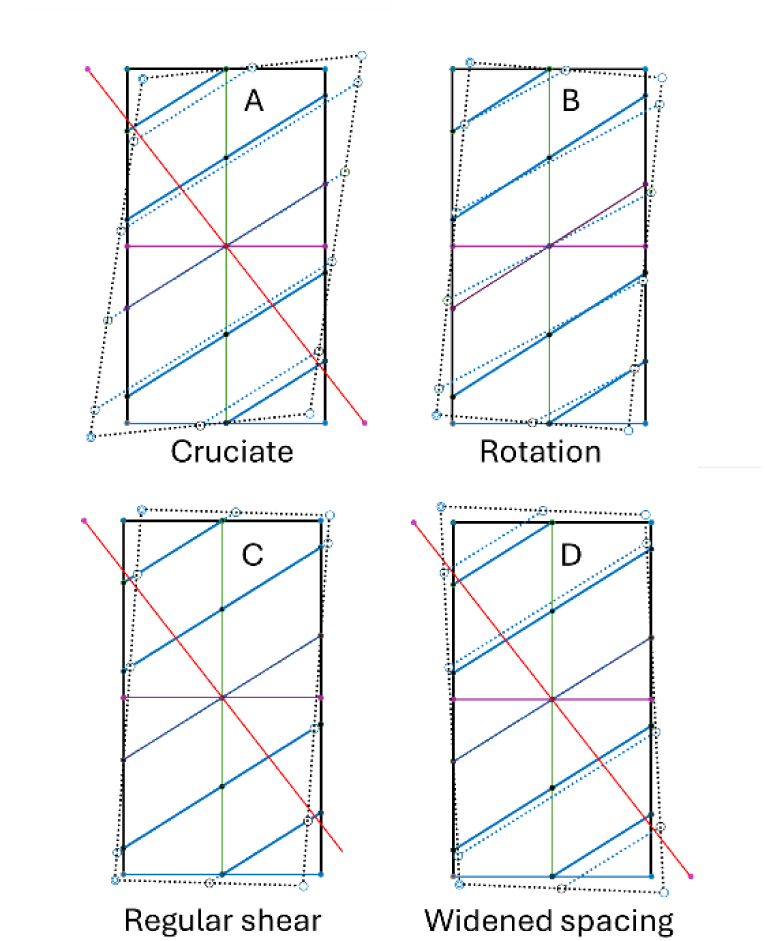
Modes of deformation of a rectangular fibril array in which the fibrils remain straight. **A**. A *cruciate* deformation modes (e.g. microfibril stretching) where elongation along the fibril axis is accompanied by orthogonal contraction as described by the Poisson ratio P: here P = 0.4. **B**. Fibril rotation by a (clockwise) angle R. **C**. Regular shear between fibrils, where each fibril slides in the same direction over the fibril below, defining a shear angle s. **D**. Widened fibril spacing, by a fraction W.

#### 2.2.1. Sliding modes

When shear between fibrils was evaluated by Keckes et al. [18], it was assumed to follow the normal engineering description: that is, successive microfibrils are displaced in the same direction by a fixed distance relative to their spacing, and thus a constant shear angle s can be defined. Here, that sliding process is called *regular shear*.

However, the sliding of successive fibrils need not be identical and thus need not conform to a constant shear angle. During the elongation of wood cell-walls with very low microfibril angle, under the rather high stresses where elastic stretching of cellulose is observed, cellulose stretching (from X-ray diffraction, XRD) did not account for all the macroscopic extension [14]. Rotation was negligible and the existence of a novel kind of fibril sliding was therefore inferred [14]. XRD data on primary cell walls at very large elongations allow the same inference [27]. In the simplest representation of this second mechanism, the direction of sliding alternates between successive fibrils. Therefore, this deformation mode is here called *interdigitated sliding*.

It can be assumed that regular shear and interdigitated sliding are simply the extremes of a continuum of sliding deformations, and that the direction of sliding might be intermediate or random rather than uniform as in regular shear or strictly alternating as in interdigitated sliding. In primary cell-walls, both sliding modes are restricted to short junction zones randomly distributed in the fibril network. Separating these two forms of sliding is maybe therefore artificial, but it simplifies both the understanding and the geometric analysis of the phenomenon. Both forms of sliding may occur between microfibrils within junction zones of primary walls or between aggregates of microfibrils, as presumed in wood [14].

An alternative way to express the existence of a continuum of sliding modes would be to define a parameter r [0.5 < r < 1] representing the proportion of sliding that is unidirectional towards the end of the fibrils nearest the apical end of the cell (right over left, RL in Fig. 1: the opposite direction is left over right, LR). Then r = RL/(LR+RL) = 1 signifies regular shear and r = 0.5 signifies interdigitated sliding, or random sliding averaging to the same magnitude as interdigitated sliding.

It is normal engineering practice to assume that what is here called regular shear is not accompanied by any lateral contraction [18], a convention followed in this study, consistent with constant volume as in solid materials. Note, however, that cell-wall volume is not necessarily constant: water can be removed or supplied from within the cell if the water activity (determined partly by pectins [28]) changes. The density of confined water may also change [29].

It can reasonably be suggested that for interdigitated sliding, lateral (Poisson) contraction accompanies elongation along the fibril axis, but the Poisson ratio is difficult to estimate. At very small (probably elastic) interdigitated sliding deformations, the data of [30] suggest that microfibril spacings should increase slightly, i.e. the Poisson ratio should be negative. For larger deformations where a stick-slip mechanism comes into play [30], gaps closing behind withdrawn fibril ends would lead to a positive Poisson ratio.

Interdigitated sliding is an example of what is here called a *cruciate* deformation mode, since elongation along the fibril axis is accompanied by cross-wise Poisson contraction. Regular shear is not.

#### 2.2.2. Fibril stretching

Elastic elongation of microfibrils has been observed by XRD and vibrational spectroscopy in wood with low microfibril angle (<10°) [14] and in axially oriented microfibrils of primary cell walls under external tension [27]. Microfibril stretching conforms to the cruciate model of deformation as described above (Fig. 2). Crystallographic Poisson ratios of approximately 0.5 have been observed for wood cellulose [31] and for intact wood [32], although curiously there seems to be no change in centre-to-centre spacing observed by SANS [33].

**Fig. 2.**
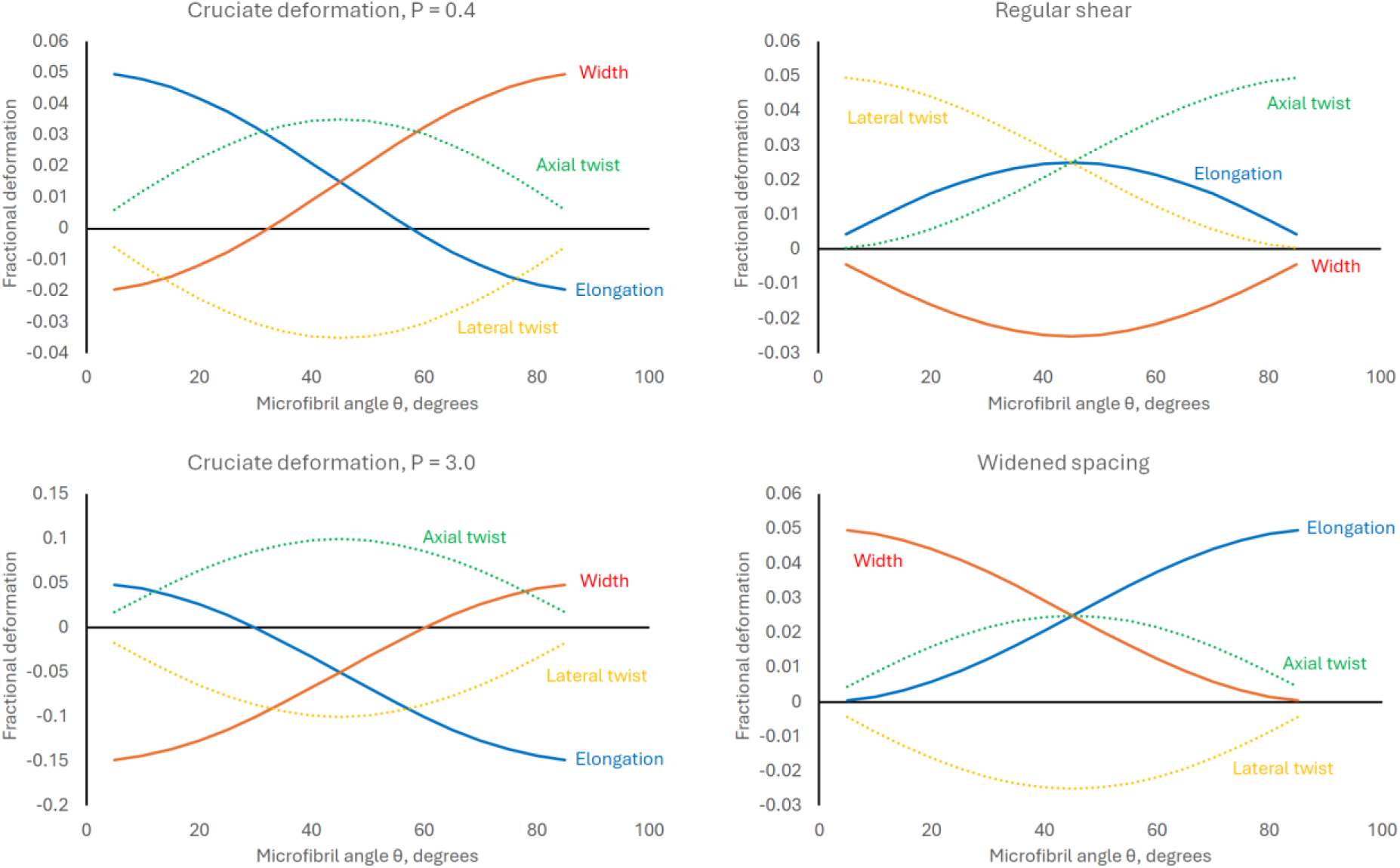
Relationship of changes in domain dimensions to microfibril angle for cruciate deformations with Poisson ratio 0.4 and 3.0, and for widened fibril spacing and regular shear. A Poisson ratio of 0.4 is potentially representative of microfibril stretching or interdigitated shear. A Poisson ratio of 3.0 is representative of the straightening of waves (2.4).

#### 2.2.3. Fibril reorientation

Fibril reorientation been observed by XRD [14], vibrational spectroscopy [34] and atomic force microscopy (AFM) [27], both in wood and in primary cell walls, where it accompanies both growth and extension under externally applied stress [1]. Where rotation counteracts twist from other deformation modes it gives an increase in both length and width. In principle rotation can be in either direction. However in practice, rotation towards the direction of stress or growth seems normally to be observed: for example, the increasingly axial microfibril orientation towards the outermost and oldest layers of an elongated primary cell wall [35].

#### 2.2.4. Widened fibril spacing

Increased spacing of microfibrils in primary cell-walls has not often been observed but was suggested by Marga et al. [19]. While respacing of wood microfibrils within macrofibrils is readily observed by small-angle neutron scattering (SANS) [33], increases in macrofibril spacing in wood would be difficult to detect except by nanoscale imaging.

It is difficult to comment on the ease or difficulty of fibril respacing without being able to specify the interaction potential between fibrils. It has been suggested that a Lennard-Jones type of potential function governs microfibril spacings in primary cell-walls, with attractive terms and a repulsive term describing contact [36]. The structure of water inserting itself between two cellulose surfaces should contribute to such a function [29, 30]. So should the nature of the cellulosic surfaces, any associated negatively charged pectic chains and any xyloglucan chains that find room for insertion [1, 15, 28]. However, much remains to be experimentally demonstrated about these phenomena, even the exact location of the relevant polymers.

#### 2.2.5. Wave formation and straightening

Wave formation in laterally oriented microfibrils has been observed for onion epidermal walls under severe external tension [27]. Coarse-grained modelling studies of primary cell-walls [21] predict that waves will form passively when transversely oriented microfibrils are compressed along the microfibril axis by narrowing of other cell-wall layers. Alternatively, forces applied perpendicular to the microfibril axis at isolated points or junction zones along each microfibril could also lead to the formation of waves, with crests and troughs at the points of application of force. These two mechanisms could occur together when a rectangular domain with high microfibril angle is stretched, and would not be easy to distinguish. Nor could either of these passive mechanisms for wave formation be easily distinguished from any waves that might result directly from sinuous paths of cellulose synthase complexess across the cell membrane. Similar sinuosity, but with only low amplitudes, contributes to the spread of fibril orientations in wood cell-walls [14]. Concerted waved patterns of large groups of microfibrils have often been observed by EM at the inner faces of primary cell-walls [19, 20, 37], although there is some doubt as to their origin. They could be a consequence of the relaxation of turgor stress, or due to shrinkage on drying prior to imaging.

When waves in axially oriented fibrils are initially present in a primary cell-wall under external tension, they tend to be pulled straight [15, 21, 27], as predicted from coarse-grained modelling [15, 21]. It is not clear if that such waves are available to be straightened during growth.

### 2.3. Geometry of individual deformations

For each deformation mode the functions for elongation, change in width and axial and lateral twist of the rectangular domain (Table 1) were derived as described in the Methods and in SI Appendix A. The basis of the derivations can be examined visually in Fig. S1.

**Table 1.**
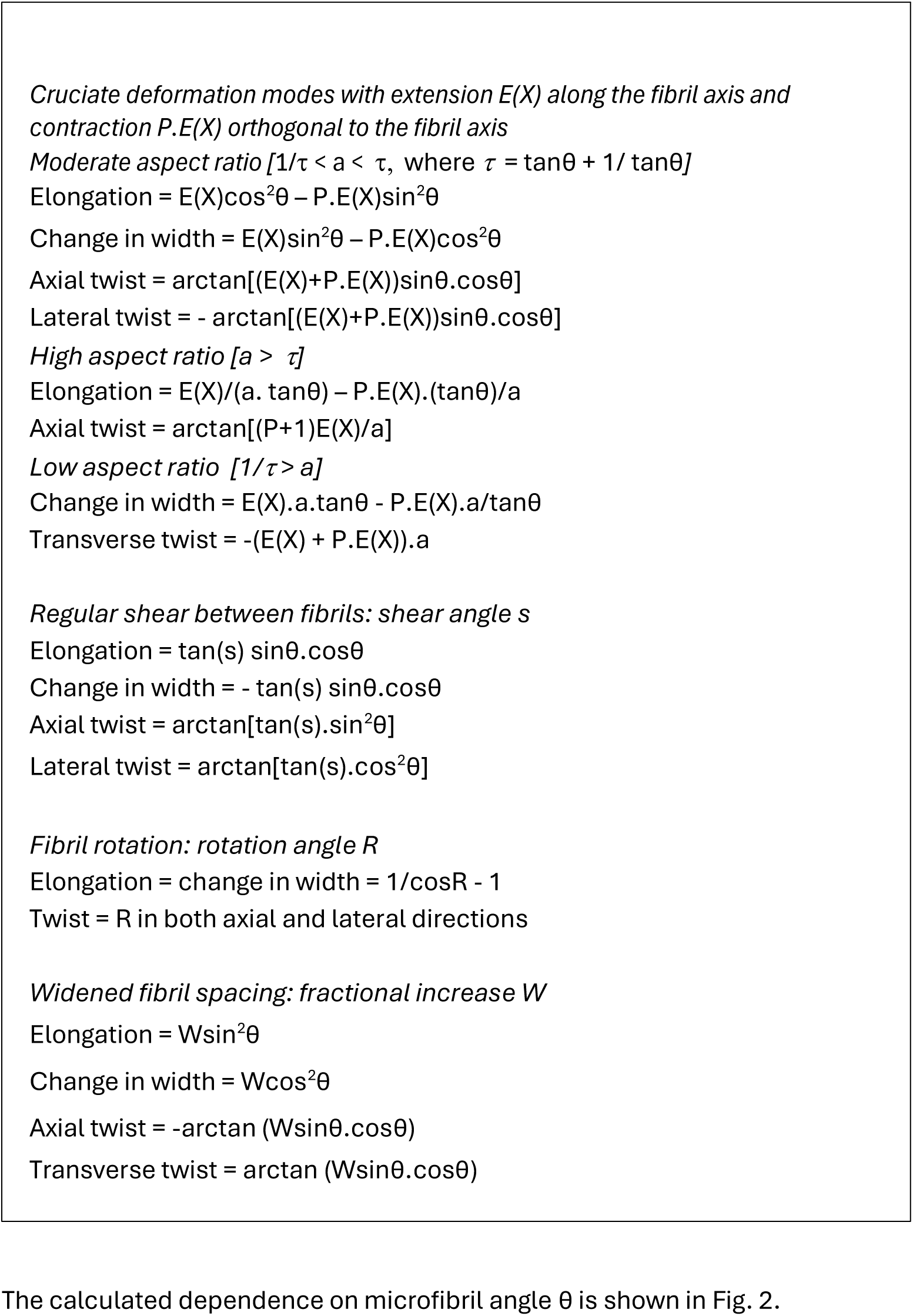
Functions used for the deformation modes.

Different functions apply to very long or very short rectangles, with aspect ratio a outside the range [1/τ < a < τ] where τ = (tanθ + 1/tanθ) (SI Appendix 1; Figs 3, S2 and S3). Examples of rectangular cell-wall domains with high aspect ratio include the edge regions of elongated primary cell walls [23]; collenchyma cell walls [38]; and the tangential walls of softwood tracheids under axial tension [14]. If the microfibril angle is close to 0° or 90°, however, these domains may still be within the range [1/τ < a < τ] (Fig. S2). That is the origin of the discontinuous behaviour of the elongation and other functions for rectangles of high or low aspect ratio.

**Fig. 3.**
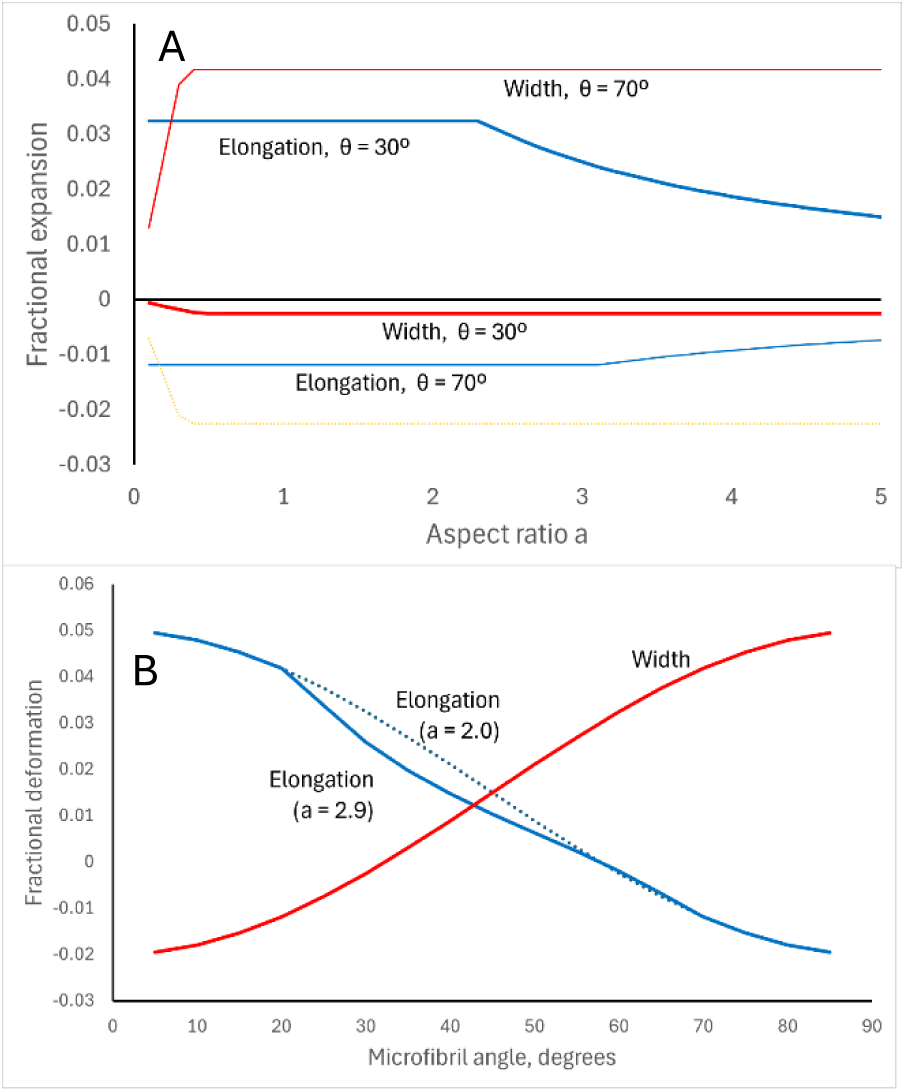
Discontinuous variation of elongation and change in width on cruciate deformation (P=0.4) of (A) rectangular domains with varying aspect ratio, at microfibril angles of 30° and 70°. (B) Variation with microfibril angle at aspect ratio a = 2.0 (moderate, dotted) or 2.9, which is sufficient to enter the high aspect ratio regime for elongation at microfibril angles much greater or less than 45°. At high aspect ratios there is no effect on width. Data on twist are in Fig. S3.

### 2.4. Geometry of wave formation and straightening

A single fibril wave can be approximated as a sinewave with amplitude 2A in Cartesian coordinates where X = distance along the axis of the wave, oriented at microfibril angle = θ to the cell axis. It is assumed that when the wave straightens, θ remains constant, i.e. the local strain is along the fibril axis not the cell axis. There is support for this view from in experiments on wood [6] and in coarse-grained (CG) modelling of primary walls [15]. Wave formation is simply the reverse of straightening.

Straightening a wave leads to elongation E(X) along the microfibril axis. The dependence of E(X) on the wave amplitude is derived in SI Appendix B and is non-linear (Figs. S4 and S5), approximated as E(X) = 0.1982A^2^ + 0.0201A.

The associated lateral contraction on wave straightening depends on how the fibrils are arranged or bundled. If separate microfibrils are sinuous and randomly arranged, with the same overall orientation but with the waves varying in amplitude and out of phase (Fig. S6), the lateral contraction on straightening can approach the mean amplitude of the waves. In contrast, if adjacent microfibrils are waved in-phase and laterally associated, the lateral contraction on straightening may be small. In intermediate cases the lateral contraction is a function of the amplitude A and the *coherence length* C of the wave pattern perpendicular to the fibril axis. For individually sinuous microfibrils C is negligibly small. For concerted wave patterns C is larger, and can reach the same order of magnitude as the wavelength [37]. The array then contracts laterally on straightening from approximately (C+A) to C, and the fractional narrowing P.E(X) = A/(C+A).

The straightening of waves, concerted or not, is an example of a cruciate deformation and can be described by the functions mentioned above, with Poisson ratios higher than in other cruciate deformation modes (Fig. S7). The geometry of wave formation, or of an increase in wave amplitude, is then a close approximation to the geometry of widened fibril spacing but with the axes transposed. When P>>1, if waves are created by local forces orthogonal to the fibrils, at junction zones corresponding to the peaks and troughs of the resulting waves, these forces will exert a large leveraged amplification (mechanical advantage) = P on the force of contraction along the fibril axis.

At the nanoscale, wave formation and straightening entail relatively small rearrangements of local structure, and have been simulated to occur before sliding deformations [21]. There is a small amount of sliding or respacing due to the curvature of concerted waves (SI Appendix 2).

### 2.5. Combinations of deformation modes giving 1% elongation with varying microfibril angle and domain width

The approach taken was to specify how the shape of the cell-scale rectangular domain changes with a small specified elongation, e.g. 1%, and then determine what combination of nanoscale deformations is required to lead to these new domain dimensions. The changes in the dimensions and shape of the initially rectangular domain are described by four specified parameters: length, width, axial twist and lateral twist (Table 1). However, there are seven unknowns, i.e. the magnitude of each of the seven nanoscale deformation modes, so a general solution is not possible. In practice the problem can be simplified because not all the deformation modes are known to occur in all circumstances (see Methods), reducing the number of unknowns to four: regular shear, interdigitated sliding, fibril respacing and fibril rotation. At very low ϑ, fibril stretching can accompany interdigitated sliding, sharing the same geometry and using the same functions. At high ϑ, low-amplitude wave formation can accompany fibril respacing, sharing similar geometry (2.8).

Solutions were then possible at all values of the microfibril angle ϑ. When axial and lateral twist were set to zero, as in most expanding plant cells embedded in multicellular tissues, the calculated magnitudes of the four deformation modes (Fig. S10) and their contributions to 1% elongation (Fig. 4) were coordinated. Their dependence on microfibril angle is shown (Fig. 4) for four representative scenarios giving different decreases or increases in relative cell-wall width (macroscopic Poisson ratio, Π). Regular shear made its largest contribution at microfibril angles close to 45°. At high microfibril angles elongation was dominated by increased mean spacing of the fibrils, arising from respacing of parallel fibrils and potentially also from wave formation. At low microfibril angles interdigitated sliding predominated, potentially accompanied at high tensile stress by fibril stretching.

**Fig. 4.**
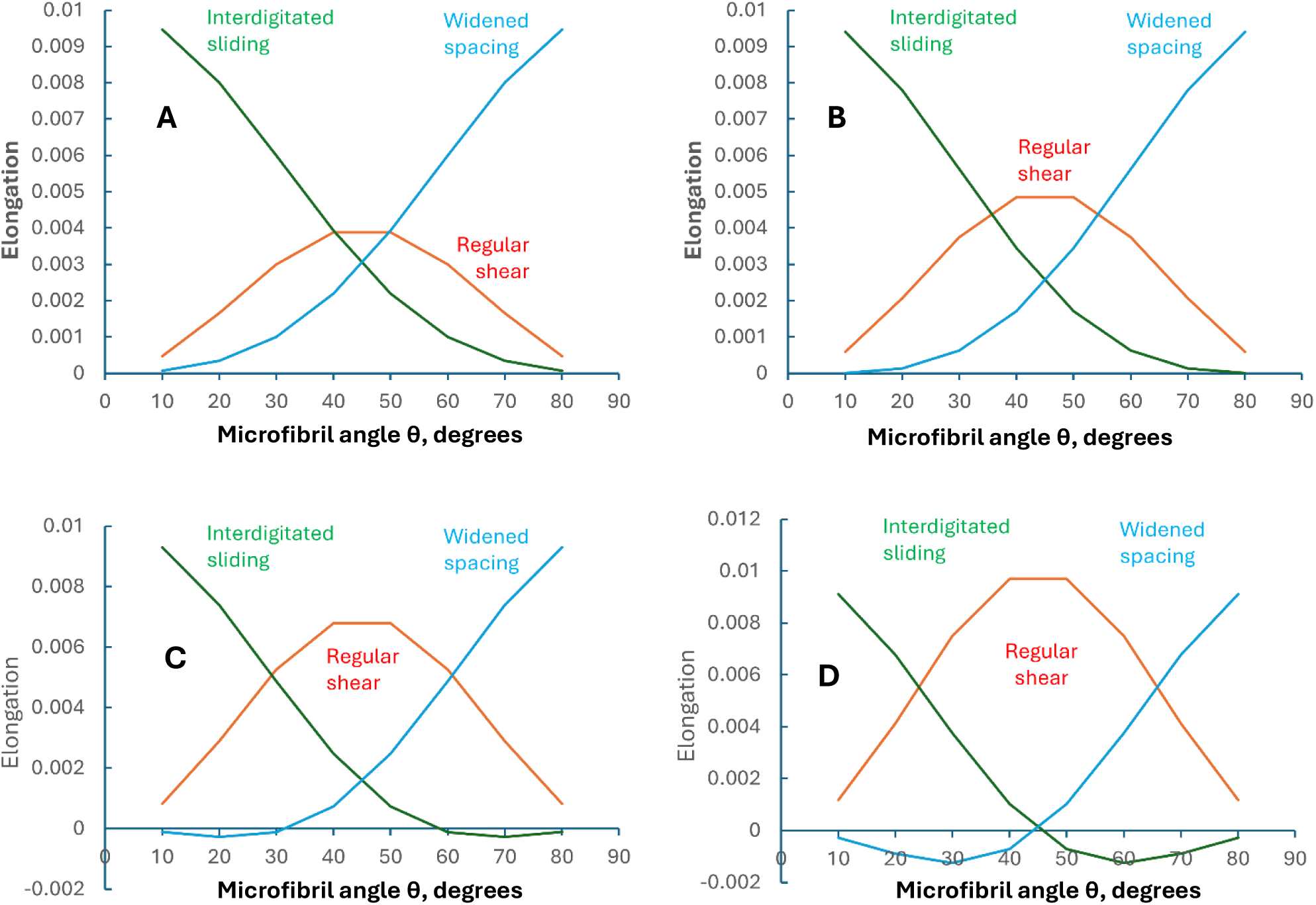
Contributions of regular shear, interdigitated sliding and widened spacing of fibrils to a total 1% elongation of a rectangular cell-wall domain, as dependent on the microfibril angle ϑ, with different widths of the domain as expressed by the macroscopic Poisson ratio Π. The contribution of fibril rotation (not shown) was small. A: macroscopic Poisson ratio Π = -0.2 (slight expansion in width as the domain elongates, representing elongating plant organs increasing in diameter). B. Π = 0, (representing parallel growth). C: Π = 0.4, (moderate narrowing, representing wood under uniaxial tension). D: Π = 1.0, (considerable narrowing, representing primary cell walls under uniaxial tension).

Increasing width of the rectangular domain (decreasing macroscopic Poisson ratio, Π) was associated with an increasing contribution of regular shear, relative to the other deformation modes (compare A to D in Fig. 4 and Fig. S10). This relationship between domain width and regular shear is shown more explicitly in Fig. S11, and was linear with Π under conditions of zero twist.

### 2.6. Incorporating twist

Macroscopic twist could in principle arise from either axial twist of cell-wall domains or from both axial and lateral twist. It was possible to constrain the above simulations to generate axial twist alone, by including large variations in the relative magnitudes of respacing, regular shear and interdigitated sliding that were nearly linear with the twist angle (Fig. S12). However outside the range 0-1° twist it was necessary to include nanoscale deformations giving increasingly negative as well as positive contributions to elongation (Fig. S13): thus one deformation mode was working against another. That outcome seems qualitatively unfavourable, and generation of axial twist alone therefore seems unlikely.

In contrast, equal, simultaneous axial and lateral twist were obtained by combinations of deformation modes that were similar, although smaller (with the exception of rotation), compared with zero twist. Most of the twist (Fig. 5) and some of the elongation (Fig. S14) were contributed by fibril rotation in the direction of twist; i.e. the microfibril angle increased to give positive twist. That scenario would be energetically more economical, as fibril rotation incurs no nanoscale rearrangement of fibril associations.

**Fig. 5.**
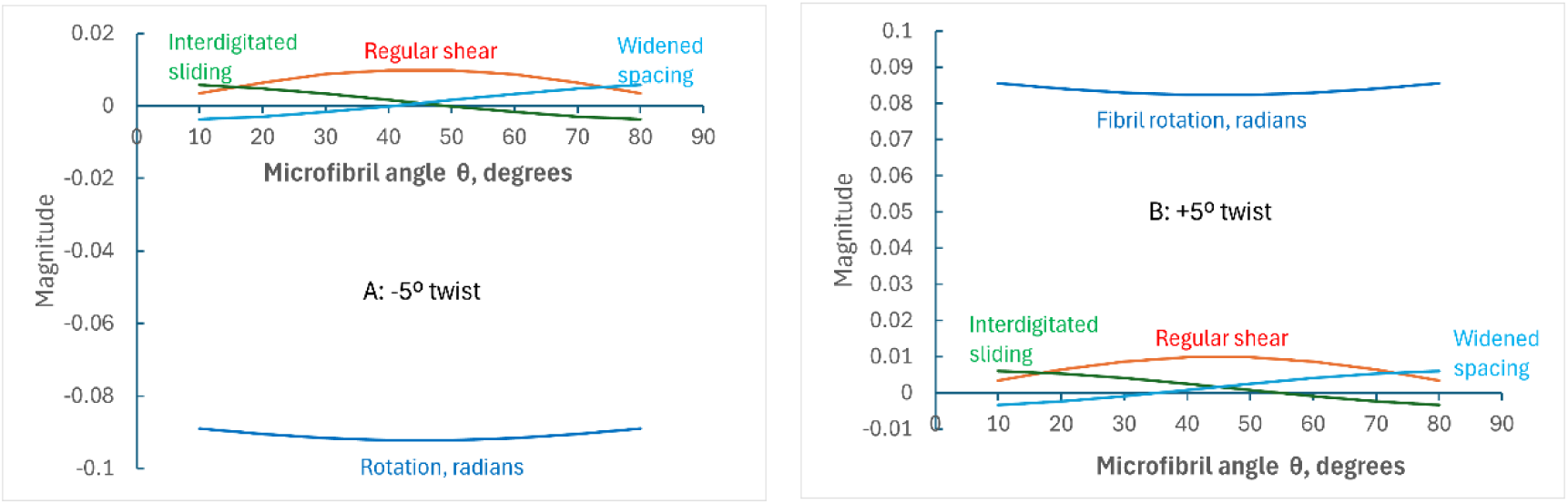
Magnitudes of deformation modes required for 1% elongation with **A:** 5° anticlockwise (-ve) axial and lateral twist and **B:** 5° clockwise (+ve) axial and lateral twist.

### 2.7. Large deformations

While elongations of 1% are commonly found in wood, primary cell walls can reach much larger elongations with evolving changes in structure that are not well described in a single simulation. An iterative approach was therefore adopted. Over 20 iterations of 5% elongation the length more than doubled and, with a macroscopic Poisson ratio of zero representing parallel growth, the microfibril angle ϑ decreased from the initial 60° to 33° (Fig. S15). The relative magnitudes of the nanoscale deformation modes changed considerably as elongation progressed (Fig. 6) in a concerted way that resulted from the constraints of zero twist and constant width.

**Fig. 6.**
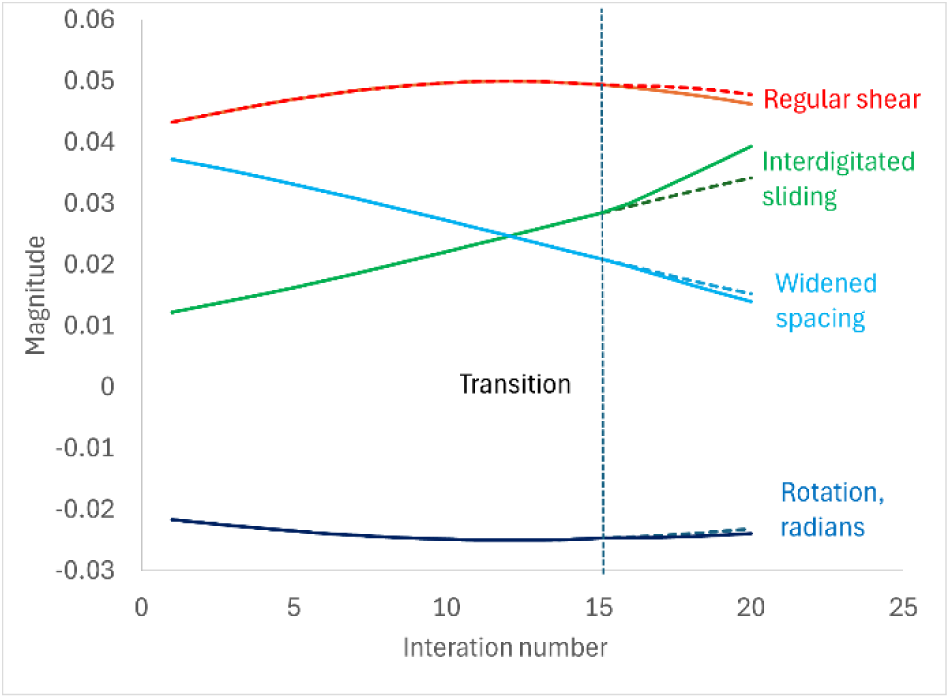
Relative magnitudes of nanoscale deformation modes during progressive iterations each of 5% elongation, with macroscopic Poisson ratio Π = 0 (i.e. no narrowing) and a starting microfibril angle of 60°. Beyond the transition point between iterations 14 and 15 where the elongation functions change, the dotted lines show the continuation of the original trend for comparison with the new, high aspect ratio regime (solid lines).

After 14 iterations the increasing aspect ratio a passed the parameter τ (= tanϑ + 1/tanϑ) (Fig. S15), marking the transition where the elongation functions change. After the transition point the magnitude of interdigitated sliding needed for 5% elongation diverged upwards with only small changes in the other deformation modes (Fig. 6), consistent with the prediction that elongation needs more nanoscale sliding, and thus potentially more energy input, in the high aspect ratio regime. There was no break in the downward slope of microfibril angle at the transition point (Fig. S15).

When the starting microfibril angle was 30° instead of 60°, values of τ were higher and remained above the aspect ratio throughout (Fig. S16), so there was no transition in the elongation functions.

A scenario with macroscopic Poisson ratio Π = 1.0 represents a primary cell-wall domain narrowing considerably as it elongates under uniaxial external tension [27]. In this scenario, because the aspect ratio rose steeply the transition to the ‘high’ aspect ratio regime occurred earlier than at Π = 0 (Fig. S17). The magnitude of the interdigitated sliding mode diverged markedly after the transition, whereas the other deformation modes did not (Fig. 7).

**Fig. 7.**
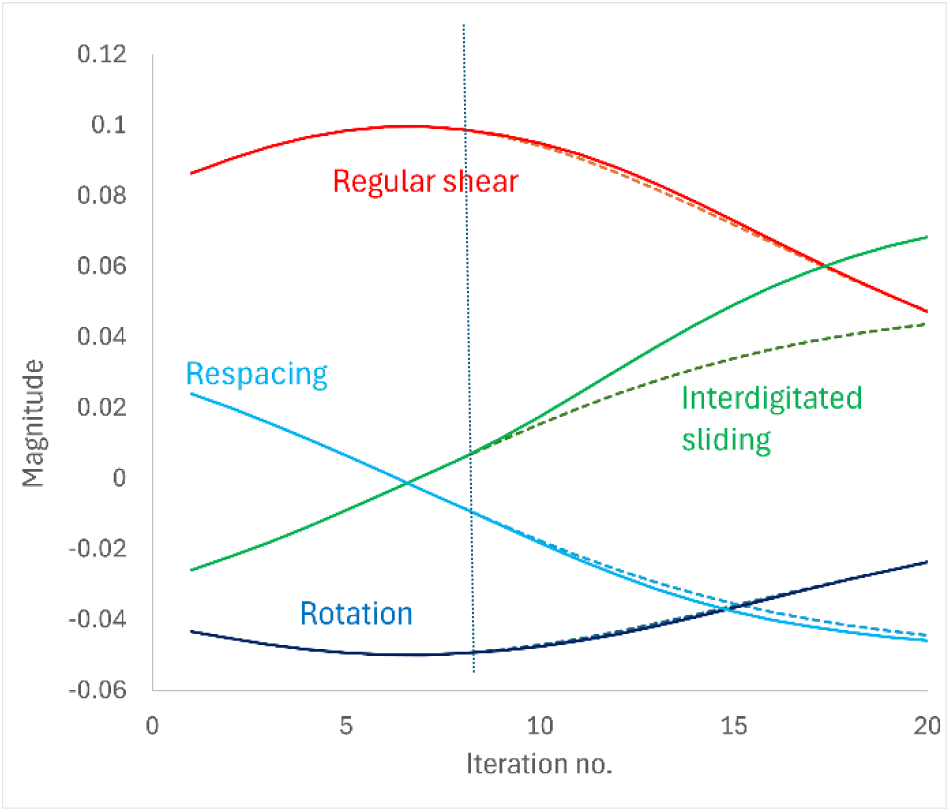
Relative magnitudes of nanoscale deformation modes during progressive iterations each of 5% elongation, with macroscopic Poisson ratio Π = 1.0 (i.e. considerable narrowing) and a starting microfibril angle of 60°. Beyond the transition point at iteration 8, the dotted lines show the continuation of the original trend, for comparison with the new, high aspect ratio regime (solid lines).

The macroscopic Poisson ratio Π need not remain constant during elongation as assumed here, although the experimental data on uniaxially stretched onion cell walls [27] suggested that Π did not vary greatly. In wood, macroscopic Poisson ratios do vary with ϑ [32], but ϑ decreases much less than in primary cell-walls at the smaller elongations observed.

### 2.8. Incorporating wave formation and straightening

It has been suggested that in primary-wall layers with axially oriented fibrils under uniaxial tension, any fibril waves that are initially present will straighten [27] before elongation occurs by combinations of nanoscale deformations that include fibril sliding, because sliding at fibril interfaces needs more structural disruption than wave straightening. Fig. 8 illustrates this scenario.

**Fig. 8.**
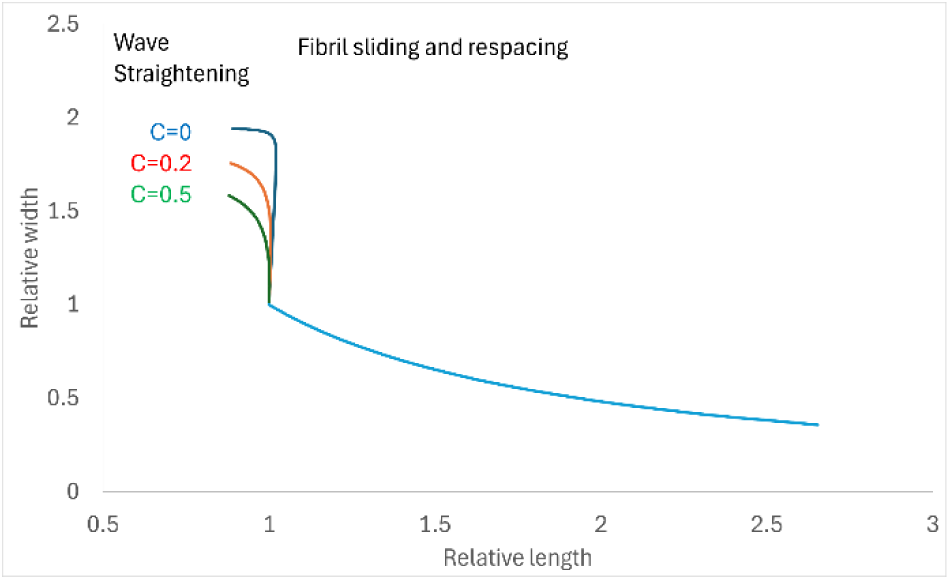
Change in width of a rectangular cell-wall domain with microfibril angle ϑ = 10° during straightening of waves with initial amplitude 0.8, followed by fibril sliding and respacing. Waves varied in coherence length C. Wave straightening was followed by 20 iterations of 5% elongation by combined respacing, regular shear and interdigitated sliding with macroscopic Poisson ratio Π = 1.0. The microfibril angle ϑ remained constant at 10° during the wave straightening phase and decreased to 1.2° by iteration 20. Due to the low value of ϑ the τ parameter remained above the aspect ratio throughout.

When the fibril waves were still of moderate amplitude, their straightening led to elongation of the cell-wall domain, presumably with low energy requirement. When the wave amplitude approached zero, the straightening of the waves provided little further elongation, and the width of the domain decreased abruptly. Indeed with non-coherent waves (C=0), there was a slight overshoot due to the projection of P.E(X) onto the domain axis x, even at ϑ = 10°. Wave straightening would probably have ceased in practice by then.

Conversely, it has been suggested that wave formation can result from lateral compression when a cell-wall lamella with high microfibril angle is forced to narrow by the lateral contraction of other lamellae [27]. Wave formation in these circumstances is compared in Fig. 9 with the narrowing effect of combined fibril sliding and respacing

**Fig. 9.**
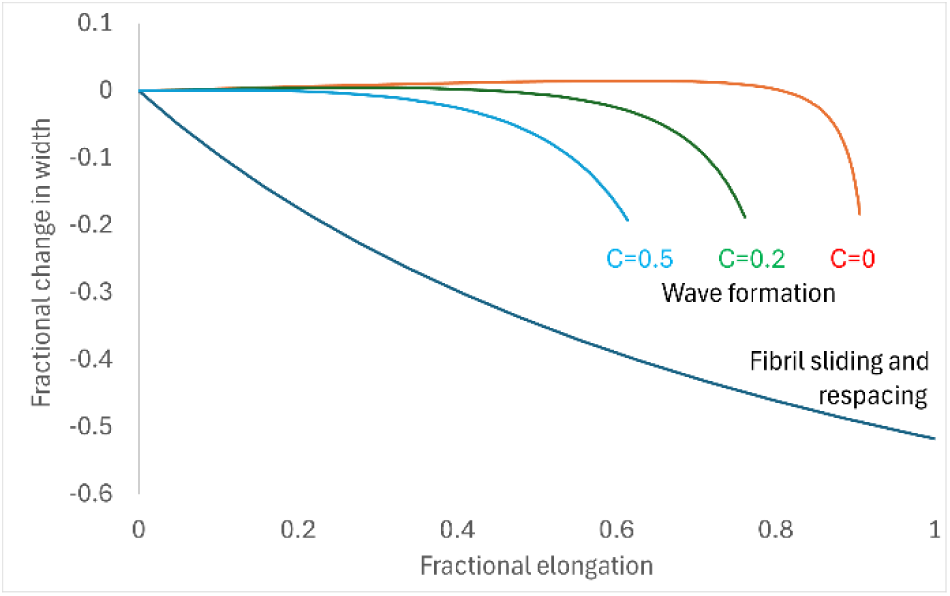
Contraction in width of a rectangular cell-wall domain with transverse fibril orientation (ϑ = 80°) by non-coherent (C=0) or coherent (C=0.2, C=0.5) wave formation. For comparison, the width contraction on equivalent elongation by 15 successive iterations of fibril sliding and respacing is also shown.

Fig. 9 shows that compared to fibril sliding and respacing, wave formation at this high microfibril angle required much more elongation of the cell-wall domain to yield any significant contraction in domain width. Indeed, for non-coherent wave patterns (C=0), there was a slight expansion in width due to the projected wave amplitude before contraction began. In these circumstances, with onely small changes in length along the fibril axis but much larger orthogonal expansion with wave amplitude, there is a close resemblance between wave formation and widened fibril spacing (with the deformation axes transposed) and wave formation can partially substitute for fibril respacing in the combined deformations (Fig. S18).

## 3. Discussion

### 3.1. Limitations

This analysis was strictly limited in its objectives, particularly through the absence of any quantitative attribution of forces, moduli or reversibility of local deformation, and through the restriction to a single lamella of a primary cell-wall, <10 nm thick, or to the S2 lamella of a wood cell-wall, some hundreds of fibril layers (several μm) thick. Deformation modes were assumed to combine additively, although non-additive interactions are also possible: for example sliding deformations synergised by fibril respacing [39] or dependent non-linearly on non-coherent wave formation [27].

Reaching a solution to the geometric equations implies that no additional stress is left within the cell-wall structure. That is an approximation: e.g. growth stresses are measurable in wood [40]; interlamellar stresses could remain after crossed-lamellate primary cell-walls elongate [5], and it is not clear if stresses at the edges of elongating cells [23] are fully dissipated.

The simulations considered here should therefore not be regarded as quantitative predictions, but merely as illustrations of how different kinds of deformation can combine during cell-wall expansion in a range of typical situations.

### 3.2. Types of fibril sliding and their control

A key finding was the need to propose two kinds of sliding deformation between fibrils, differentiating between what are here called *regular shear* and *interdigitated sliding.* These sliding motions are recognised to represent the two extremes of a continuum. Regular shear and interdigitated sliding can both contribute to the elongation of a rectangular domain, but have different relationships to width and twist.

Width is a *constraint* set to match the observed behaviour of an experimental system; whereas in a growing, cross-laminate cell-wall zero twist is a *restraint* mechanically imposed by attachment between successive lamellae and between walls of adjacent cells. Under zero-twist restraint, specifying the width constraint (macroscopic Poisson ratio, Π) determines the relative elongation contributions of all four deformation modes shown in Fig. 4, readjusting when the microfoibril angle decreases at large elongations. Anything that changes one deformation mode will force changes in the rest (Fig. S11) and in width (Π).

There are implications for cross-lamellate cells undergoing diffuse elongation growth [41, 42]. Increasing regular shear increases Π (Fig. S11). During growth, sliding between microfibrils at specific junction zones called ‘hot spots’ is facilitated by expansins [43]. However, it is not clear what is distinctive about hotspot structure, nor whether, under biaxial turgor stress, expansin action would discriminate in favour of the RL sliding required for regular shear.

Under uniaxial tension the local force vectors for interdigitated sliding, regular shear and widened spacing are proportionate to cosϑ, cos(2ϑ) and sinϑ respectively, maximal at ϑ = 0°, 45° and 90°. Perhaps fortuitously, these maxima for the local force vectors correspond to the maximal contributions of the four deformation modes to elongation, as shown in Fig. 4, even though the calculations leading to Fig. 4 take no account of force. Away from the maxima the correspondence is not exact: the elongation contributions decrease more steeply. But it seems that each deformation mode is preferentially deployed at microfibril angles where its driving force is maximal.

In wood under tensile stress, enzymatically facilitated sliding is absent but otherwise, similar considerations may determine how contributions of deformation modes vary with microfibril angle and how the macroscopic Poisson ratio emerges [14]. Sliding is presumably between macrofibrils or larger units, not microfibrils as in primary cell-walls. Curled, tapered strands observed at tensile fracture surfaces in wood [44] may imply locked-in shear between units larger than macrofibrils.

### 3.3. Large deformations and the influence of aspect ratio

An unforeseen outcome was the discontinuous dependence of elongation, width and twist on the aspect ratio of the rectangular domain (Fig. 3). Discontinuous dimensional functions are also a feature of other shapes, such as a stretched hexagon [27], that could correspond to one wall of a plant cell.

The microfibril angle decreased during large elongations and the relative magnitudes of the nanoscale deformation modes needed to change accordingly, controlled by the width constraint and the zero-twist restraint. Figs 6 and 7 show that as a primary-wall lamella with a typical microfibril angle of 60° elongated, especially if it also narrowed under uniaxial tension, the aspect ratio of the domain could cross the transition at a = τ where the elongation functions change. The main nanoscale consequence was an increase in the magnitude of interdigitated shear (Figs 6 and 7). It is noticeable that many of the cell facets in simple, elongating plant tissues fall within the ‘moderate’ range of aspect ratios, where elongation is maximal relative to the required magnitude of the local sliding deformations.

At higher and lower initial microfibril angles, τ remained large enough to prevent the transition point from being reached, even at large elongations (Figs S2 and S16). This would apply to the long, narrow cell-wall domains at cell edges where the microfibril angle remains close to 90° [45], and to wood cell walls where the microfibril angle is small. Other very long cells, such as cotton hairs, show tip growth [46] or are discontinuous with respect to microfibril angle [22].

### 3.4. Waves and changes in fibril spacing

In primary cell-walls with laterally oriented fibrils (high microfibril angle) wave formation may contribute to the elongation and narrowing of the cell-wall (SI Appendix B). Cell-wall elongation increases linearly with wave amplitude, steeply for non-coherent waves. Cell-wall narrowing too increases with wave amplitude but to a negligible extent until the amplitude is quite large, [e.g. A = 0.2). Thus wave formation is not effectively driven by narrowing of one cell-wall lamellae forced by other lamellae of the same cell-wall, as suggested [27]. Waves are more likely to form in these circumstances by the alternative mechanism where external tensile stress is transmitted to discontinuous junction zones that become the peaks and troughs of the wave pattern.

Whatever forces drive their formation, waves with low amplitude and low coherence are very similar (with axes transposed) to the widened spacing of straight fibrils with respect to the geometric behaviour simulated here, although distinguishable experimentally by imaging [47] or diffraction [27]. Only respacing of straight fibrils is illustrated in Fig. 1 but both respacing and wave formation can in principle contribute to the increased mean spacing of microfibrils that is the main contributor to elongation at high microfibril angles (SI Appendix B, Fig. S18).

The geometric similarity between respacing and wave formation might resolve a paradox. It is widely observed that plant cells grow preferentially at right angles to the predominant cellulose orientation (e.g. [48, 49] but - intuitively and from Fig. 4 - this implies increased mean spacing of the cellulose fibrils. Respacing of parallel fibrils does occur [19], but is often thought to be less important than fibril sliding [41]. Possibly, low-amplitude waves contribute to the increase in mean fibril spacing in these circumstances.

In contrast, images of the ‘edge’ domains of elongating primary cell-walls [45] suggest that their microfibrils remain straight and parallel at approximately ϑ = 90°, implying that respacing of straight fibrils, not wave formation, permits elongation of these domains.

Parallel, axially oriented microfibrils with widened spacing have been observed at large elongations of primary cell-walls under uniaxial tension [27]. If the oldest, outermost lamellae of growing cells [50] have similarly widened spacing of their axial fibrils, with fewer remaining junction zones to resist interdigitated sliding, it might explain the long-standing puzzle that these oldest lamellae fail to prevent elongation growth [35].

### 3.5. Twist

Twist at organ level requires twist at cellular level [51] and is observed in, for example: tendrils of climbing plants [51, 52], the twisted growth of certain mutants [53] and spiral grain in trees [54, 55], from which the ensuing twist in kiln-dried wood is a major economic problem.

While organ-level twist could in principle arise from axial twist alone at cellular level, Figs 5 and S12-S14 suggest that axial and lateral twist are more likely to occur together. as implicitly envisaged for wood [56]. In the primary walls of elongating cells, some deviation from a regular alternating cross-lamellate wall structure seems necessary [53].

### 3.6. Conclusion

This study was limited to cell-wall domains of simple rectangular shape, stretched or growing in a simple uniform way. The geometry-based approach was more limited than, e.g., coarse-grained modelling [21] or continuum mechanics [57] but was complementary and unexpectedly productive. It demonstrates the principle that the shape of growing cells can derive from appropriate combinations of nanoscale deformations within the primary cell wall, constrained purely by geometry.

More complex changes in cell shape may emerge from non-uniform expansion of cell walls, such as those of the leaf epidermis [58], hair primordia [59] or conifer tracheids with pit fields in their radial walls [44]. There, the balance of deformation modes needs also to be non-uniform across the cell surface and would be better simulated using an array of circular micro-domains (SI Appendix C).

## 4. Methods

The derivation of the functions describing the geometry of each deformation mode is described in SI Appendix A and illustrated in Fig. S1. A key step was the correct definition of the normal axis for each component of each deformation mode, where deformation is zero. The deformation functions that were derived are shown in Table 1.

For combinations of deformation modes, the approach taken was to specify how the shape of the cell-scale rectangular domain changes with a small specified elongation, e.g. 1%, and then determine what combination of nanoscale deformations is required to lead to these new domain dimensions, as described in SI Appendix D, Supplementary Methods.

## Acknowledgements

The author thanks C. Anderson and D. Cosgrove for discussions that led to the development of the ideas described here.

## Notes

### Competing Interest Statement

The authors have declared no competing interest.

### Summary of Updates

Additional material has been included to show how the deformation modes originally described can be combined together

