## Supplementary Information for "Deformation geometry of cellulose fibril arrays constraining the stretching and growth of plant cell walls"

#### **SI Appendix A.** *Derivation of cruciate, respacing and regular shear deformation functions*

In the cruciate modes of local deformation, extension E(X) in the direction X of the microfibril axis is accompanied by contraction P.E(X) in the direction perpendicular to the microfibrils [33]. The ratio of lateral contraction to axial expansion is the Poisson ratio P, as defined with respect to the microfibril axis (not, here, the cell axis, with respect to which the macroscopic Poisson ratio $\Pi$ is defined).

The first step is to define the two orthogonal neutral axes on which elongation and lateral contraction, respectively, are equal to zero. Incorrect designation of these neutral axes mixes fibril shear with the other two deformation modes. The contributions of elongation in the direction of the fibril orientation and lateral contraction are each calculated from the distance from the appropriate neutral axis (Fig. S1). For elongation in the direction of the fibril orientation, the neutral axis is the line normal to the fibrils, passing through the origin at the centre of the rectangle. For lateral contraction the neutral axis is the line parallel to the fibrils, passing through the same origin.

For rectangular domains of moderate aspect ratio a (all domains with $0.5 < a < 2$, and domains outside this range at high or low microfibril angles), the elongation of the rectangle is measured in the y direction between the midpoints of the top and bottom, and contains contributions from both E(X) in the direction of the fibrils and P.E(X) in the orthogonal direction, each multiplied by the distance of the top and bottom midpoints from the corresponding neutral axis (Fig. S1A). The axial twist is measured from the x coordinate of the same displacement. The change in the width of the rectangle and the lateral twist are calculated in an analogous way from the (x,y) displacement of the mid-points of the sides of the rectangle. (Fig. 2; SI Appendix A and Fig. S1A). Perpendiculars drawn from the neutral axes to these four mid-points form the lines along which axial elongation and lateral contraction of the rectangular domain are measured, except when they extend outside the boundary of the initial rectangle, in which case only that part of the line lying within the rectangle is measured.

Twist has the same magnitude in both axial and lateral directions but is opposite in sign. In consequence the rectangle distorts to a parallelogram (Fig. 1). That is why the dimensional changes are calculated between the midpoints of the edges. Rotation cannot compensate simultaneously for the axial and the lateral twist associated with the cruciate deformation modes.

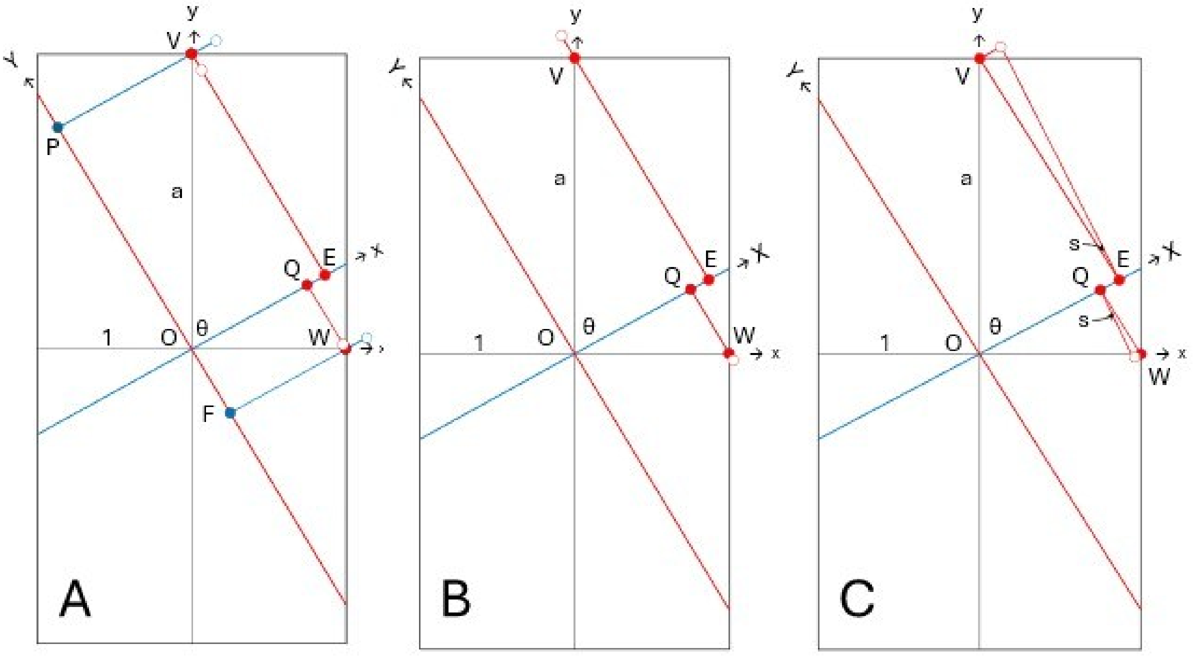


**Fig. S1.** Geometry of cruciate, respacing and regular shear deformation modes for the upper half of a rectangular fibril array with half-width unity and aspect ratio a (within the 'moderate' range). The coordinate system (x,y) for the array is related to the coordinate system for the fibril orientation (X,Y) by the microfibril angle θ, sharing origin O. The fibril orientation is shown blue and the orthogonal direction is shown red, with neutral axes X and Y passing through O. Elongation of the rectangle and axial twist are measured from change in the y and x coordinates at point V. Change in width of the rectangle and lateral twist are measured from change in the x and y coordinates at point W. Filled circles denote the initial state of the system and open circles denote the deformed state. **A**: Cruciate deformations where expansion along the fibril axis is accompanied by contraction orthogonal to the fibrils, each relative to the distance from the respective neutral axis Y or X. **B:** Respacing deformation comprising expansion orthogonal to the fibrils, from neutral axis X. **C:** Regular shear tan (s) parallel to the fibril orientation, relative to the distance from neutral axis X.

*Derivation of functions for cruciate deformations*

For cruciate deformations the elongation of the rectangle is derived from the sum of the y contributions from both E(X) and P.E(X) at point V. The axial twist is derived from the sum of the x contributions from both E(X) and P.E(X) at point V. The change in width of the rectangle is derived from the sum of the x contributions from both E(X) and P.E(X) at point W. The lateral twist is derived from the sum of the y contributions from both E(X) and P.E(X) at point W.

The (x,y) contribution from E(X) at point V is given by E(X).[PV]. The distance [PV] = [OV].cos θ = a. cos θ .

The fibril elongation from E(X), $\delta X = E(X).a.\cos\theta$

with y projection $\delta y = E(X)\ a.\cos^2\theta$ and x projection $\delta x = E(X)\ a\cos\theta.\sin\theta$.

The (x,y) contribution from P.E(X) at point V is given by P.E(X).[EV]. The distance [EV] = [OV].sin θ = a. sin θ .

The orthogonal contraction from P.E(X), $\delta X = P.E(X).a.\sin\theta$

with y projection $\delta y = P.E(X)\ a.\sin^2\theta$ and x projection $\delta x = -P.E(X)\ a\cos\theta.\sin\theta$.

Summing the contributions from E(X) and P.E(X):

Fractional elongation $\delta y/a = E(X)\cos^2\theta - P.E(X)\sin^2\theta$

Tan axial twist $\delta x/a = (1+P)E(X)\cos\theta\sin\theta$

The change in width and lateral twist are calculated similarly:

The (x,y) contribution from E(X) at point W is given by E(X).[FW]. The distance [FW] = [OW].sin θ = sin θ .

The fibril elongation from E(X), $\delta X = E(X).\sin\theta$

with y projection $\delta y = E(X)\cos\theta.\sin\theta$ and x projection $\delta x = E(X)\sin^2\theta$

The (x,y) contribution from P.E(X) at point W is given by P.E(X).[QW]. The distance [QW] = [OW].cos θ = cos θ .

The orthogonal contraction from P.E(X), $\delta X = P.E(X).\cos\theta$

with y projection $\delta y = P.E(X)\cos\theta.\sin\theta$ and x projection $\delta x = -P.E(X)\cos^2\theta$.

Summing the contributions from E(X) and P.E(X):

Fractional change in width $\delta x/1 = E(X)\sin^2\theta - P.E(X)\cos^2\theta$

Tan lateral twist $-\delta y/1 = -(1+P)E(X)\cos\theta\sin\theta$

In rectangular domains of high aspect ratio, points P and E lie outwith the sides of the rectangle and the dimensions PV and EV are replaced by the distance from the left and right sides respectively. Thus PV = a. cos θ is replaced by 1/sin θ and EV = a. sin θ is replaced by 1/cos θ. Then Elongation = $E(X)/(a.\tan\theta) – P.E(X).(\tan\theta)/a$ and Axial twist = $\mathrm{atan}[(P+1)E(X)/a]$.

Similarly, when the aspect ratio is low, points Q and E lie outwith the top and bottom of the rectangle and the dimensions PV and EV are replaced by the distance from these boundaries.

*Derivation of functions for respacing deformation*

Widened spacing deformation is derived similarly to the cruciate deformations but making use of Fig. S1B. The expansion E(X) along the microfibril axis X is zero and the orthogonal contraction P.E(X) is replaced by the expansion W, such that -W = P.E(X).

The (x,y) expansion at point V is given by W.[EV]. The distance [EV] = [OV].sin θ = a. sin θ .

The (x,y) expansion at point V is therefore W.a.sin $\theta$

with y projection $\delta y$ = W.a. $\sin^2 \theta$ and x projection $\delta x$ = -W.a cos $\theta$. sin $\theta$.

The (x,y) expansion at point W is given by W.[QW]. The distance [QW] = [OW].cos $\theta$ = cos $\theta$ .

The (x,y) expansion at point W is therefore W.cos $\theta$

with y projection $\delta y$ = -W. cos $\theta$. sin $\theta$ and x projection $\delta x$ = W. $\cos^2 \theta$.

Then fractional elongation = W. $\sin^2 \theta$

Change in width = W. $\cos^2 \theta$

Axial twist = -W.cos $\theta$. sin $\theta$

Lateral twist = W. cos $\theta$. sin $\theta$

*Derivation of functions for regular shear deformation*

Regular shear is derived similarly to the cruciate deformations making use of Fig. S1C.

Regular shear tan(s) displaces point V by tan(s) [EV] = tan(s).a.sin $\theta$

with y projection tan(s).a. cos $\theta$.sin $\theta$ and x projection tan(s).a.$\sin^2 \theta$.

Regular shear tan(s) displaces point W by tan(s) [QW] = tan(s).cos $\theta$

with y projection -tan(s).$\cos^2 \theta$ and x projection -tan(s). cos $\theta$.sin $\theta$.

It is assumed that there is no contribution from a Poisson effect.

Then fractional elongation = tan(s).a. $\cos^2 \theta$ /a = tan(s). cos $\theta$.sin $\theta$

Change in width = -tan(s). cos $\theta$.sin $\theta$

Axial twist = tan(s).a.cos $\theta$.sin $\theta$/a = tan(s).$\sin^2 \theta$

Lateral twist = tan(s).$\cos^2 \theta$

### List of Symbols

Fractional elongation of cell wall: E

Microfibril angle (angle between cell axis and microfibril axis): $\theta$

Aspect ratio of cell-wall domain: a

Fractional elongation along fibril axis (fibril stretching, interdigitated shear, wave straightening): E(X)

Nanoscale Poisson ratio (fibril stretching, interdigitated shear): P

Fractional contraction at right angles to the fibril axis (fibril stretching, interdigitated shear): E(Y) = P.E(X)

Fractional expansion at right angles to the fibril axis (widened fibril spacing): W

Shear angle (regular shear): s

Macroscopic Poisson ratio (for the elongating rectangular domain): $\Pi$

Distance along fibril axis: X

Wave amplitude: A

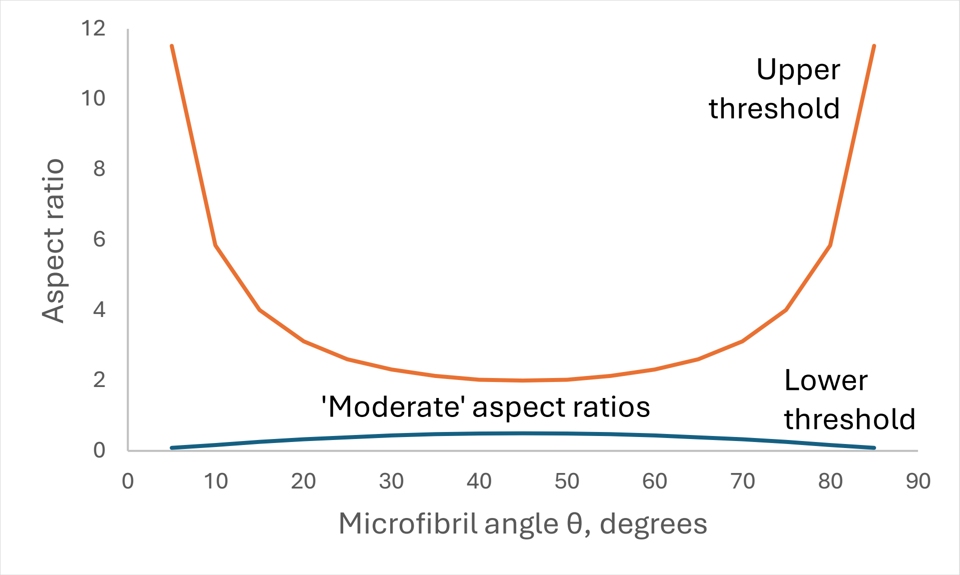


**Fig. S2.** The three ranges of aspect ratios where the functions defining cruciate deformation of a rectangular domain differ, with upper and lower thresholds for the region of 'moderate' aspect ratio.

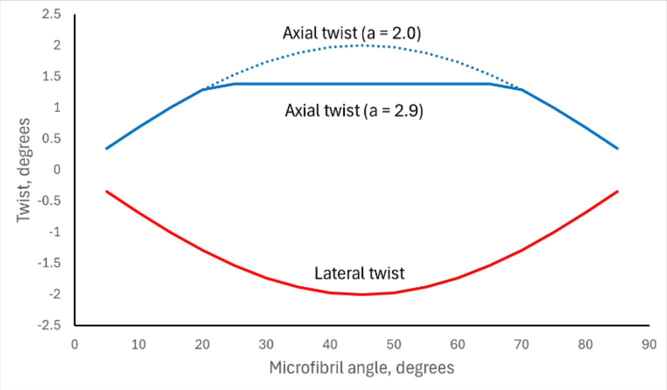


**Fig. S3.** Twist due to cruciate deformation in a rectangular domain with aspect ratio a = 2.9, where the axial twist function is in the 'moderate' aspect ratio regime when the microfibril angle θ is high or low but in the high aspect ratio regime when the microfibril angle is intermediate.

### **SI Appendix B.** *Formation and straightening of wave patterns*

First consider a single waving fibril

$Y = A.\sin(X)$ ...................................................................(1)

where the ratio of amplitude to wavelength = $A/\pi$

At any point x along the waveform the gradient

$dY/dX = A.\cos(X)$

When a segment of the fibril straightens its fractional elongation

$E(X) = (1 + A.\cos^2 X)^{0.5} - 1$ ..............................................................(2)

Integrating (2) from X=0 to X=2π gives the overall fractional elongation along the microfibril axis, shown below as a function of the amplitude A. A power-law fit to this relation gives

$E(X) = 0.1982A^2 + 0.0201A$

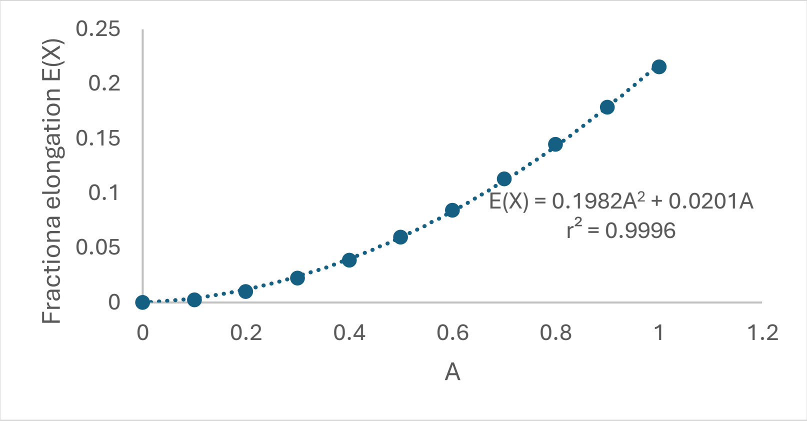


**Fig. S4.** Elongation E(X) along the fibril axis when a sine wave of amplitude A straightens fully. The relationship can be fitted with the quadratic function $E(X) = 0.1982A^2 + 0.0201A$. Wave formation is the reverse of wave straightening.

The ratio A/E(X) is the Poisson ratio P for straightening of a single wave. The Poisson ratio for wave straightening is always greater than unity and is particularly large when the wave amplitude is small (Fig. S5)

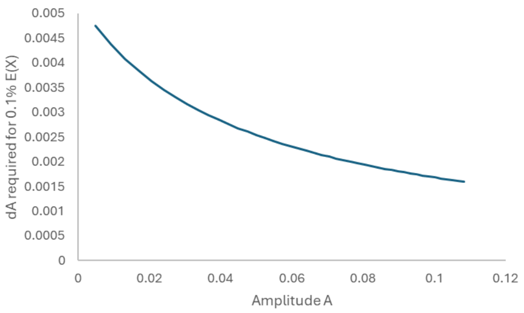


**Fig. S5** Change in amplitude A resulting in a fixed 0.1% contraction in length E(X) of fibrils with varying amplitudes of non-coherent waves.

*Non-coherent arrays of single fibrils*

Waves with varying amplitude and sporadic junction zones (Fig S6) are typically present in primary-wall domains.

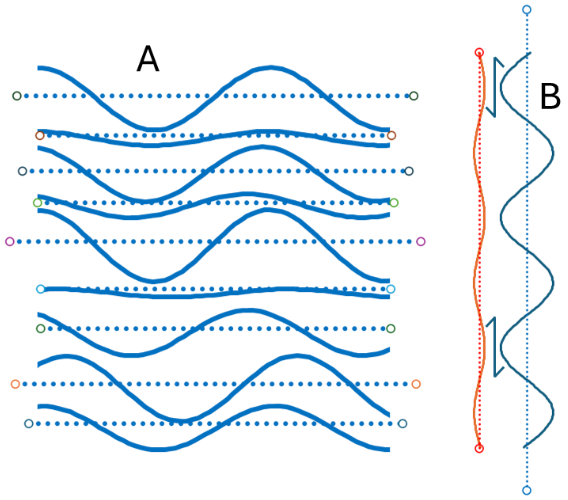


**Fig. S6. A** Straightening of an array of fibrils with non-coherent waves of varying phase and amplitude, approaching one another at junction zones. The straightened fibrils are shown as dotted lines. **B.** Differential elongation when waves of different amplitude straighten creates shear stresses at junction zones.

Straightening of the fibrils in an array like that shown in Fig. S6A is limited by those waves that are initially of lower amplitude, unless some of the junction zones deform or separate. The varying magnitude of extension means that the junction zones come under shear stress, greatest for the junction zones that link fibrils with high and low wave amplitude (Fig S6B). When the whole array extends by a fixed amount, the larger-amplitude waves do not straighten completely, and the decrease in their amplitude is less than the waves that initially were of lower amplitude.

When an array of single fibrils with high microfibril angle is forced to contract along their length by narrowing of other lamellae in the same cell-wall, waves can form in fibrils that were initially straight. Waves already present also expand in amplitude but by a smaller amount (Fig. S5), subjecting junction zones to shear stress as in Fig. S6B. Wave formation may also result when tension along the axis of the rectangle resolves into forces acting, orthogonal to the fibril axis, on inter-junction fibril segments, leading to contraction along the fibril axis as above. The contractile force has a mechanical advantage equal to the Poisson ratio for each fibril, and may be very large (Fig. S7) for fibrils that are initially almost straight. The functioning of tension wood has been suggested to involve a similar mechanism.

*Coherent and partially coherent waves*

On straightening, the contraction P.E(X) perpendicular to the microfibril axis depends on the coherence length C, as well as on the amplitude A. On straightening, a coherent domain of microfibrils will contract laterally from approx. (C+A) to C. The fractional change in width is then P.E(X) = A/(C+A). P, the apparent Poisson ratio for the straightening of coherent waves, is greater than unity but smaller than for non-coherent waves (Fig. S7).

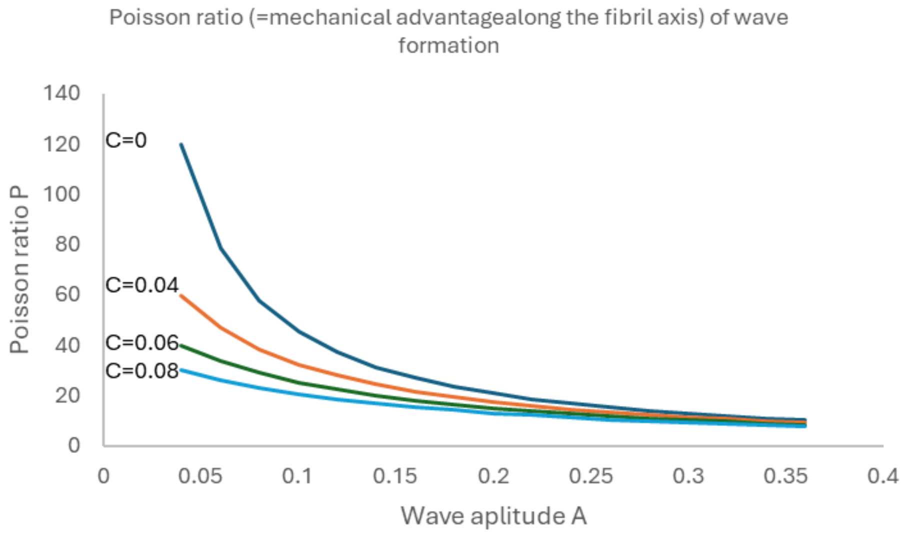


**Fig. S7.** Dependence of the Poisson ratio P on the amplitude A and coherence length C of waves in a partially coherent fibril array.

The above values of E(X) (Fig. S4) and P (Fig S7) may be substituted into the standard cruciate functions for the elongation, width change and twist of the rectangular cell-wall domain.

*Shear or respacing during coherent wave formation?*

If a coherent array of waved fibrils are simply stacked vertically, their spacing is reduced in the sloping sections according to the cosine of the local first derivative of the wave function. Alternatively, the spacing can be held constant: the radius of curvature will then increase in convex regions and decrease in concave regions, leading to shear in the sloping regions between. The changes in radius and the shear magnitude are cumulative with the coherence length C of the array. Whether it is energetically more favourable for fibrils to rearrange by shear or by respacing is in general unknown, but the cumulative nature of the shear alternative makes shear less likely than respacing when the coherence length is large.

### **SI Appendix C.** *Deformation geometry of circular domains*

The initial state of the system is described by a circular fibril array of diameter 2r with the fibrils aligned diagonally within it at an angle θ, the microfibril angle, to the cell axis. The cell axis and the axis of applied unidirectional tension are vertical. In practice, the deformed array was constructed in Cartesian coordinates with the microfibril axis vertical, and was then rotated to the microfibril angle θ (or θ - twist) using the transform (x,y) → (xcos θ + ysin θ, ycos θ - xsin θ).

The microfibril array was then deformed according to the deformation mode in question. The elongation was determined as the vertical projection of the distance between the highest and lowest points on the ellipse, and the change in width from the right and left extremities. The deformation functions for each deformation mode were checked numerically from plots of the dimensions of the original circle and the deformed ellipse.

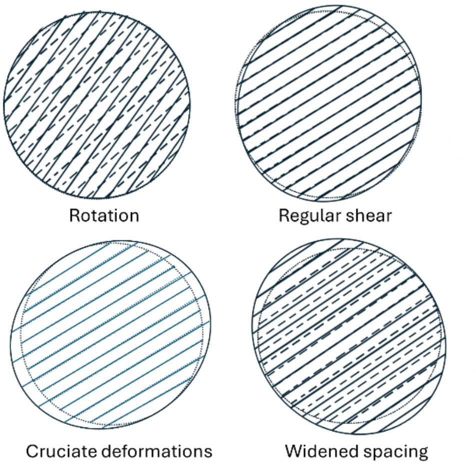


**Fig. S8** Deformation modes for circular fibril arrays.

Fig. S8 shows the principal deformation modes of initially circular domains. The functions for elongation and width change are similar but not always identical to the corresponding functions for rectangular domains. In the case of regular shear the relationship of elongation or width change to microfibril angle θ is shifted. Maximum elongation and minimum width are reached when the microfibril angle θ = approximately 45°, as expected for engineering shear. The dependence is therefore approximately on sin(2θ), but more precisely, the sine curves are slightly displaced by an increment b (-b for width) that depends linearly on the shear angle s. It was determined empirically that b = 1+s/2.89 (Fig. S9).

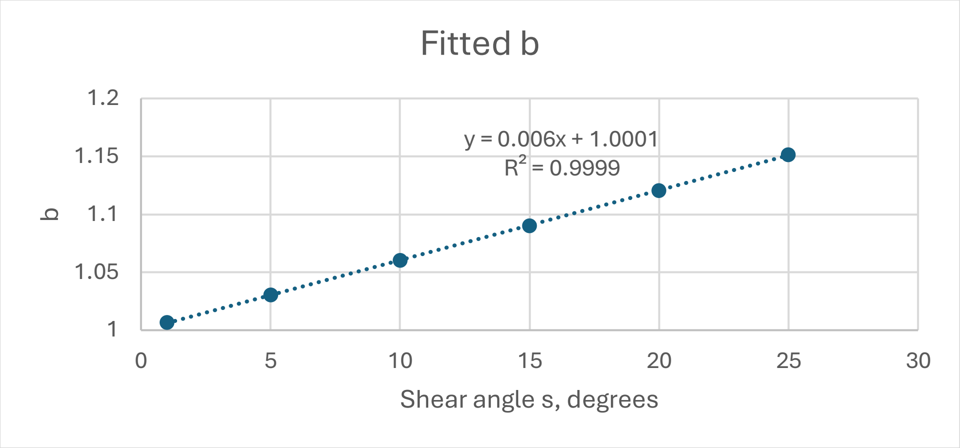


**Fig. S9.** Adjustment parameter b for the relationship between elongation and width associated with regular shear in a circular domain.

For an initially circular domain the concept of twist of the domain boundaries is meaningless. Cruciate deformations introduce ellipticity in the fibril direction while regular shear introduces ellipticity at 45º to the fibril orientation. These directions remain constant, independent of the magnitude of the deformation, so there is no twist from each of these deformation modes when considered individually. However, introducing regular shear or fibril rotation into a domain that is already elliptical due to a cruciate deformation will twist the long axis of the ellipse away from the fibril orientation by an angle dependent on the ratio of the magnitudes of the two deformation modes concerned.

Using a grid of circular domains whose deformation geometry is analysed in this way may be a practical alternative to examining an entire cell-wall facet when the cell is complex in shape.

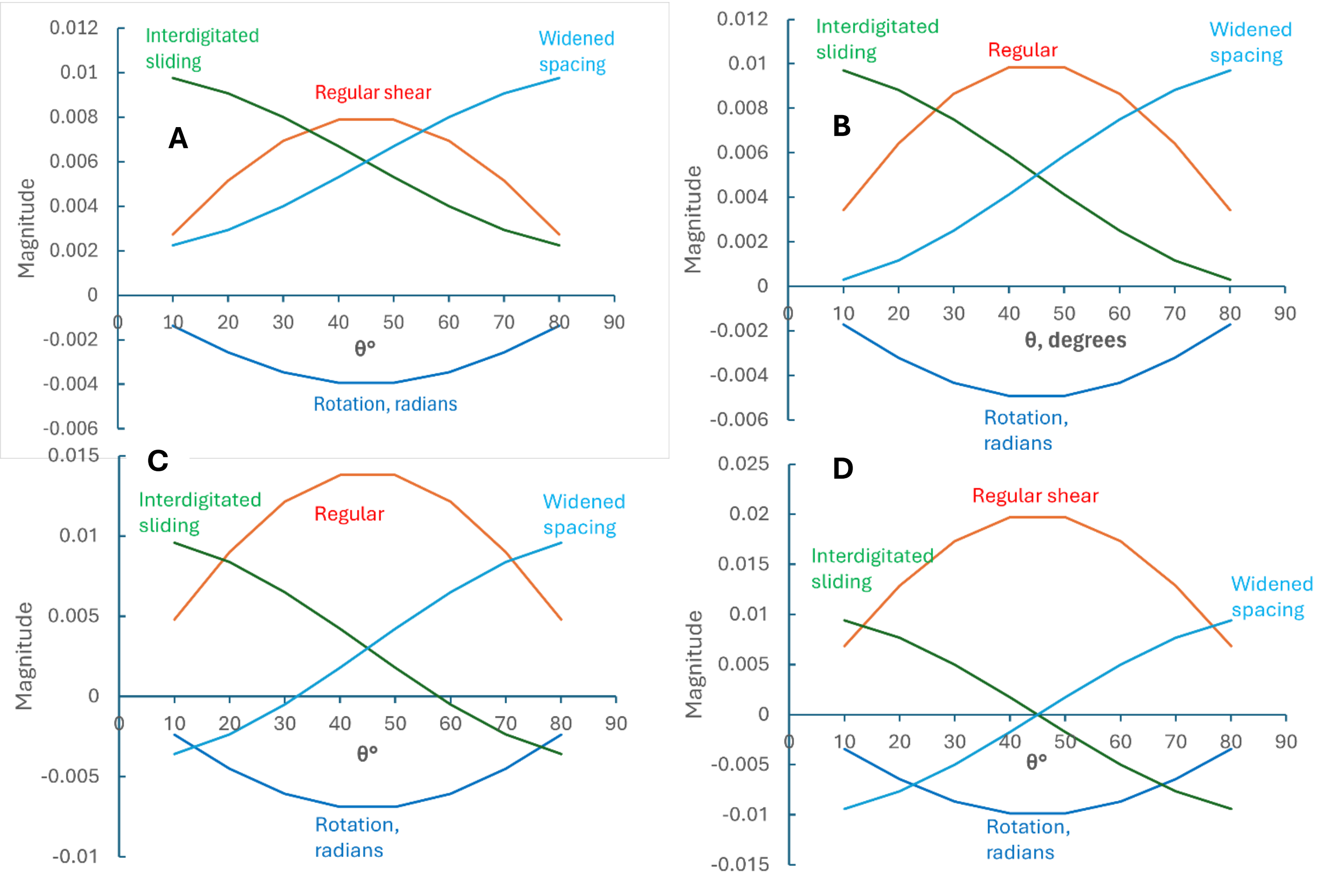


**Fig. S10.** Magnitudes of regular shear, interdigitated sliding, widened fibril spacing and fibril rotation required to give 1% elongation of a rectangular cell-wall domain with different simultaneous changes in the width of the domain, i.e. macroscopic Poisson ratios P. **A**. P = -0.2 (slight width increase as in growing cells that increase in diameter as they elongate). **B**. P = 0 (no change in width as in cells undergoing parallel elongation growth) **C**. P = 0.4 (moderate narrowing as in wood under uniaxial tension). **D**. P = 1.0 (considerable narrowing as in primary cell-walls under uniaxial tension).

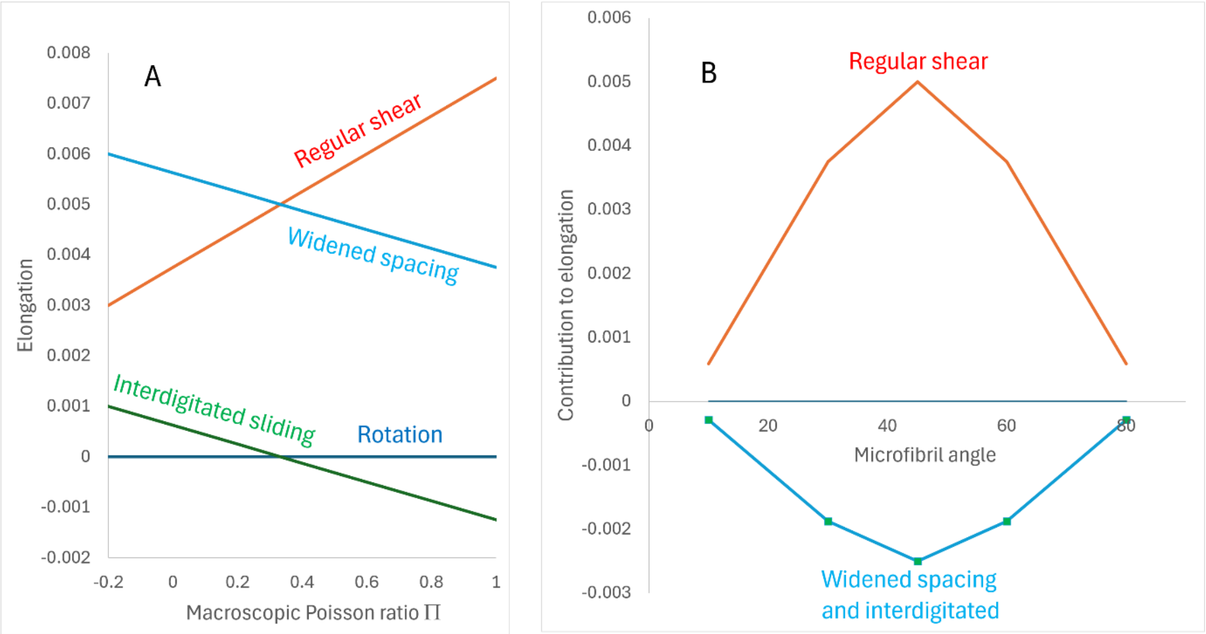


**Fig S11.** Dependence of changes in domain width with elongation (macroscopic Poisson ratio Π) on combinations of deformation modes. **A:** Linear increase in the elongation due to regular shear, and parallel decreases in the elongation due to widened spacing and interdigitated sliding, at microfibril angle θ = 60º. **B:** Microfibril angle dependence of the slopes (as in A) of variation of elongation contributed by each deformation mode, with macroscopic Poisson ratio Π. The curves for widened spacing and interdigitated sliding coincide and their ratio to regular shear is constant at -0.5. This constant relationship means that Π can be considered as being controlled by the relative contribution of regular shear.

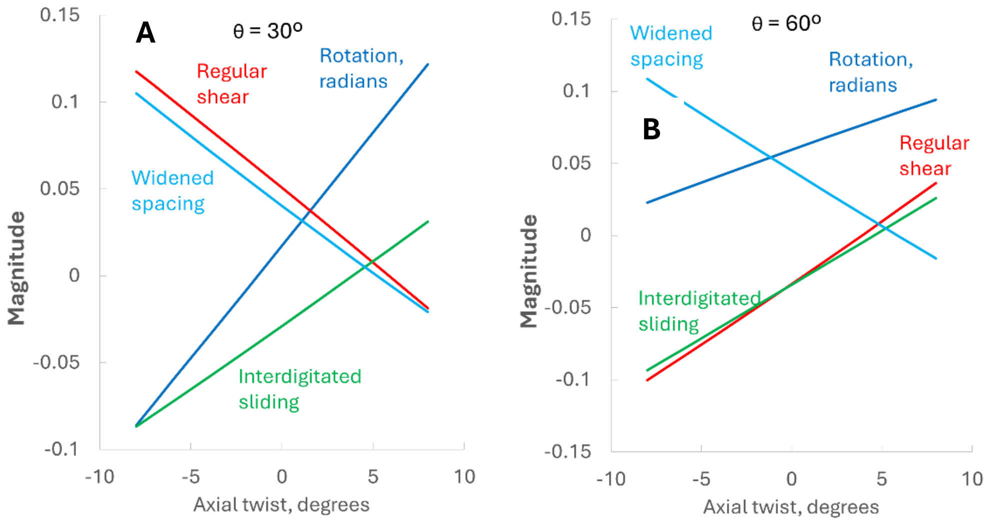


**Fig S12.** Magnitudes of regular shear, interdigitated sliding, fibril respacing and fibril rotation required to give 1% elongation of the rectangular domain, with varying degrees of axial twist and zero lateral twist. **A:** microfibril angle θ = 30º. **B:** microfibril angle θ = 60º.

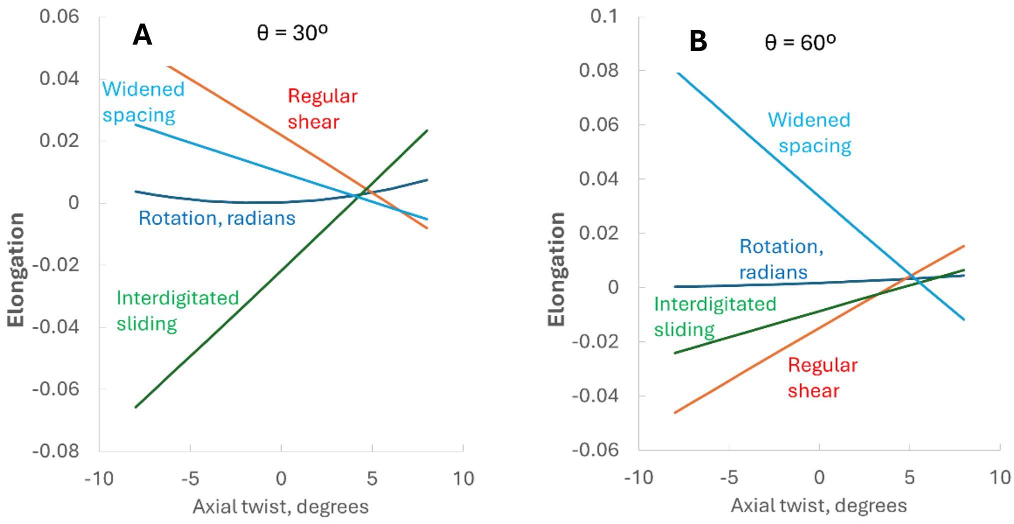


**Fig S13.** Contributions to elongation due to regular shear, interdigitated sliding, fibril respacing and fibril rotation required to give 1% elongation of the rectangular domain, with varying degrees of axial twist. **A:** microfibril angle θ = 30º. **B:** microfibril angle θ = 60º.

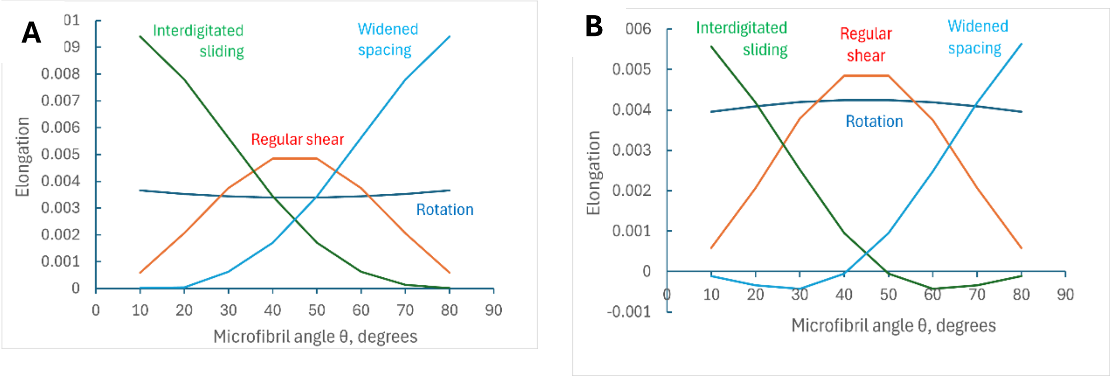


**Fig S14.** Contributions to elongation due to regular shear, interdigitated sliding, fibril respacing and fibril rotation required to give 1% elongation of the rectangular domain, with **A**: -5º axial and lateral twist (anticlockwise); B: +5º axial and lateral twist (clockwise).

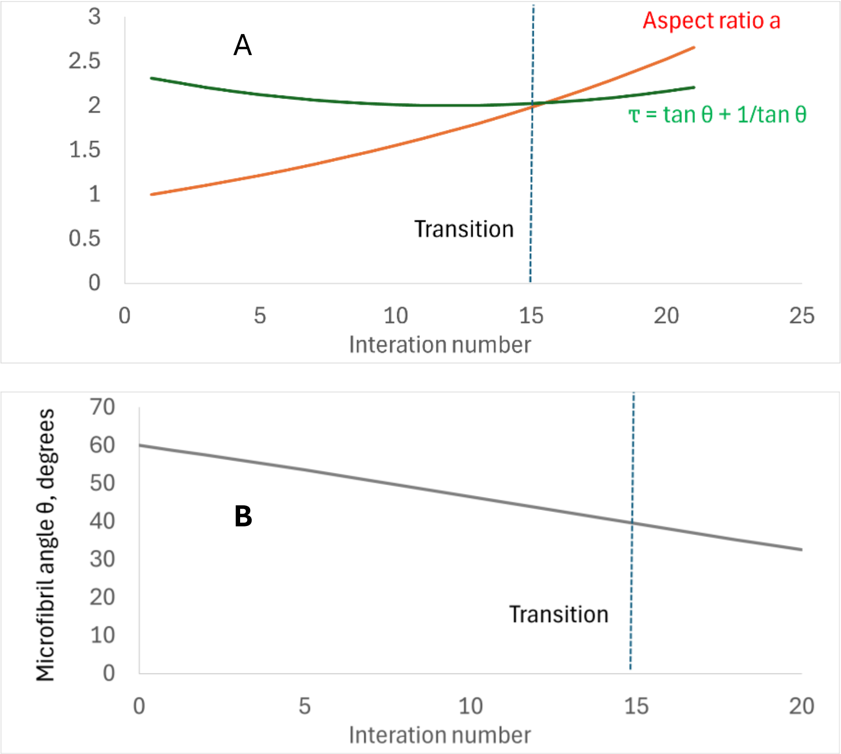


**Fig S15.** Rectangular domain with initial microfibril angle θ = 60°, during 20 iterations each of 5% elongation, P = 0. **A:** aspect ratio a and the parameter τ. The point where a and τ cross marks the transition from the elongation functions for moderate aspect ratio to the elongation functions for high aspect ratio. **B:** Decreasing microfibril angle θ with elongation.

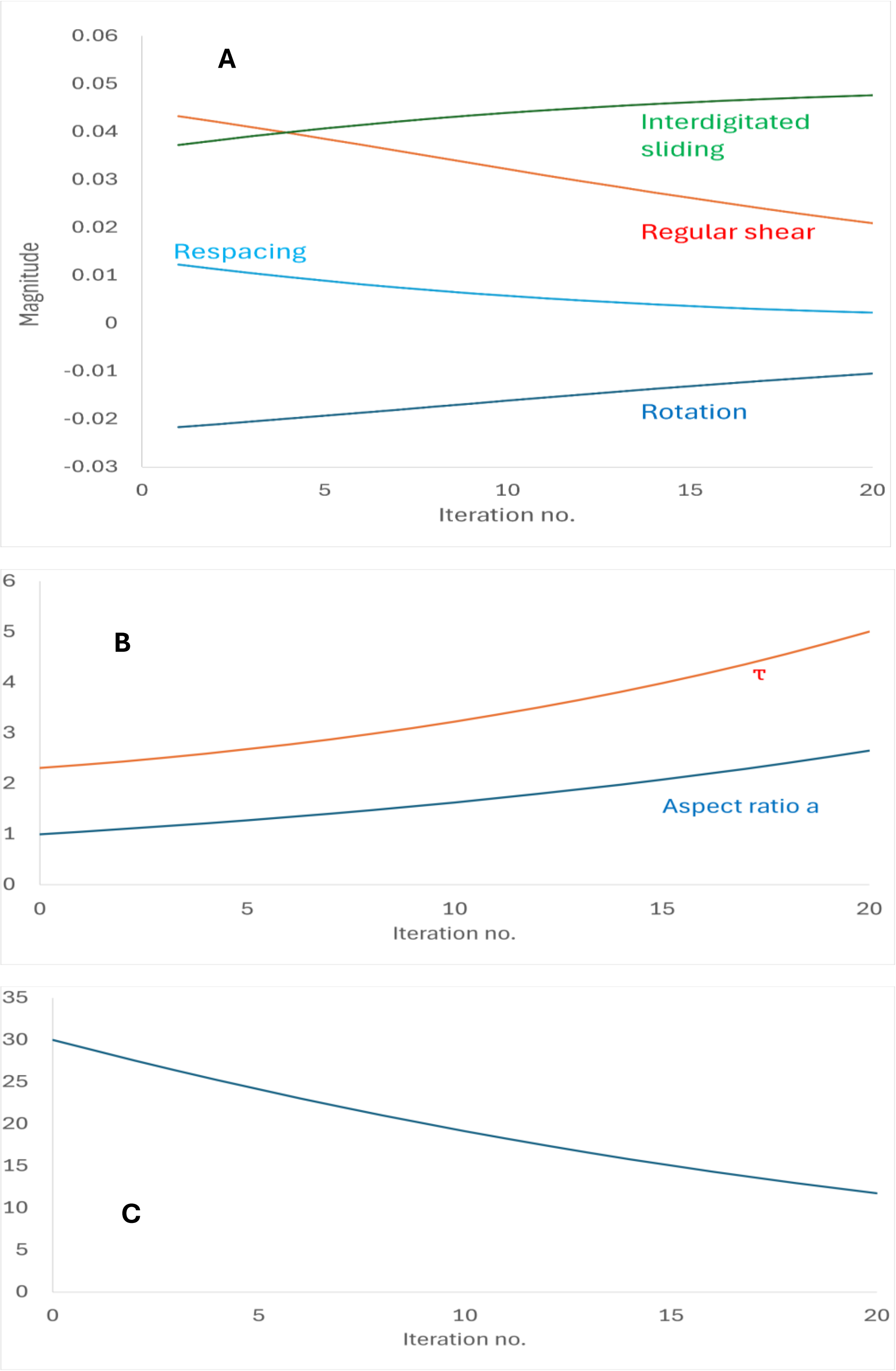


**Fig S16.** Rectangular domain with initial microfibril angle θ = 30º, during 20 iterations each of 5% elongation, P = 0. **A:** Relative magnitude of each nanoscale deformation mode. **B:** aspect ratio a and the parameter τ. The parameters a and τ do not cross, so the elongation functions for moderate aspect ratio apply throughout. **C:** Decreasing microfibril angle θ with elongation.

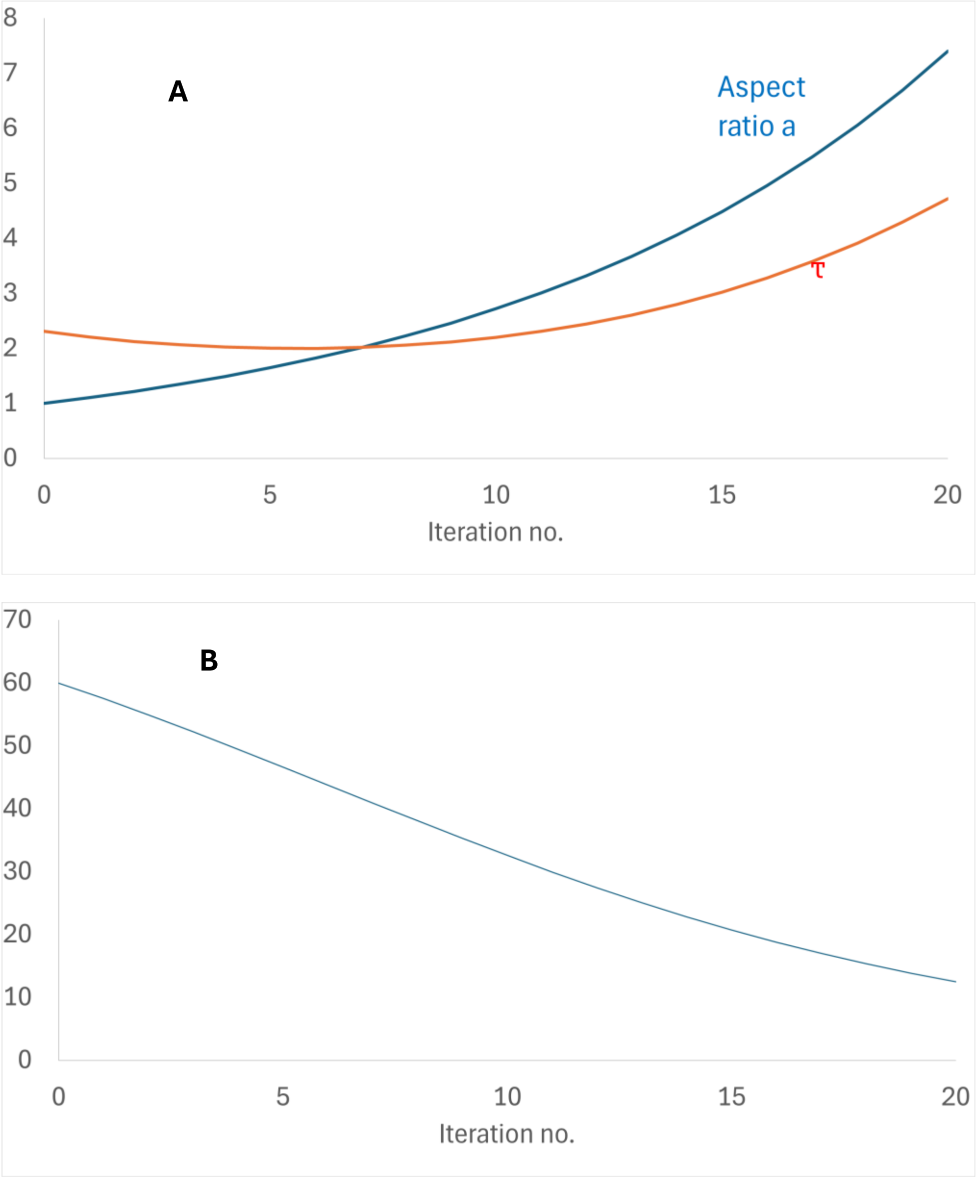


**Fig S17.** Rectangular domain with initial microfibril angle θ = 60º, during 20 iterations each of 5% elongation, P = 1.0. **A:** aspect ratio a and the parameter τ. The point where a and τ cross marks the transition from the elongation functions for moderate aspect ratio to the elongation functions for high aspect ratio. **B:** Decreasing microfibril angle θ with elongation.

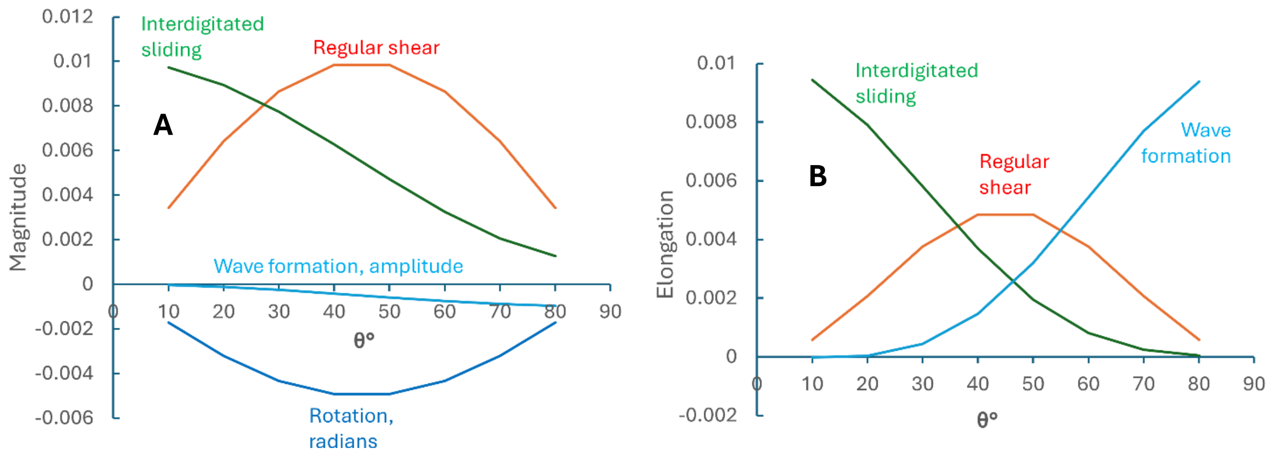


**Fig S18.** Low-amplitude wave formation substituting for respacing of parallel fibrils. **A:** Magnitudes of regular shear, interdigitated sliding, wave formation and fibril rotation required to give 1% elongation of the rectangular domain, as a function of microfibril angle θ at $\Pi = 0$. **B:** Contributions to the total elongation resulting from these individual deformations (Compare with Figure 2B) .

### SI Appendix D. *Supplementary methods*

Analysis of combinations of deformation modes was carried out as follows. From Table 1 it can be seen that the changes in the dimensions and shape of the initially rectangular domain are described by four specified parameters: length, width, axial twist and lateral twist. However, there are seven unknowns, i.e. the magnitude of each of the seven nanoscale deformation modes [40], so a general solution is not possible. In practice the problem can be simplified because not all the deformation modes are known to occur in all circumstances. Three such exceptions are noted below.

Fibril stretching is observed only when the microfibril orientation is close to axial, under high external tensile stress [15, 28]. Fibril stretching and interdigitated sliding have identical geometry. In wood with microfibril angle <10°, they occur together and microfibril stretching can be individually quantified by wide-angle X-ray or neutron scattering [15, 28]: it typically accounts for less than half of the total elongation and is elastic, whereas interdigitated sliding is plastic.

When coherent waves are initially present, their straightening requires little nanoscale disruption, is observed before other modes of deformation begin [28], and can therefore be evaluated as a separate initial deformation phase.

In cell-wall domains with very high microfibril angle [40], when elongation of the cell wall under external tension is accompanied by reduction in width [28] the nanoscale geometries of fibril respacing and the formation of non-coherent waves are difficult to distinguish [40]. In these circumstances it was initially assumed that only one or the other was present and the results were compared.

With these simplifications, the number of known variables was reduced to equality with the number of unknowns. The resulting simultaneous equations were solved numerically by least-squares optimisation as described below.

Combining deformation modes: optimising fitted parameters

1. The half-height and half-width of the rectangular domain and the microfibril angle $\theta$ were specified. The parameter ($\tan\theta + 1/\tan\theta$) was calculated from the specified $\theta$.

2. Target parameters were inserted for fractional elongation, fractional change in width, axial twist and transverse twist.

3. Arbitrary values of 0.01 were inserted for the magnitudes of the four deformation modes to be combined, normally fibril rotation, regular shear, interdigitated sliding and fibril respacing. The Poisson ratio P for interdigitated sliding (or any other cruciate deformation mode included) was specified.

4. For each deformation mode, the functions in Table 1 were used to calculate fractional elongation, fractional change in width, axial twist and transverse twist.

5. The deviations $\delta(i)$ (calculated – target) in fractional elongation, fractional change in width, axial twist and transverse twist were calculated for each deformation mode, squared and added together to give the sum of squares $\Sigma\delta$.

6. The SOLVER add-in in Excel was used to minimise $\Sigma\delta$. by varying the magnitudes of the four deformation modes as unconstrained variables. The GRG Nonlinear algorithm with forward derivatives was used and negative values of the unconstrained variables were permitted. Solutions were accepted when $\Sigma\delta$ $<10^{-12}$.
